# Orange is the new white: taxonomic revision of Antarctic *Tritonia* species (Gastropoda: Nudibranchia)

**DOI:** 10.1101/2020.07.31.230664

**Authors:** Maria Eleonora Rossi, Conxita Avila, Juan Moles

## Abstract

Among nudibranch molluscs, the family Tritoniidae gathers taxa with unclear phylogenetic position, such as some species of the genus *Tritonia* Cuvier, 1798. Currently, 35 valid species belong to this genus and only three of them are found in the Southern Ocean, namely *T. challengeriana* Bergh, 1884, *T. dantarti* Ballesteros & Avila, 2006, and *T. vorax* (Odhner, 1926). In this study, we shed light on the long-term discussed systematics and taxonomy of Antarctic *Tritonia* species using morpho-anatomical and molecular techniques. Samples from the Weddell Sea and Bouvet Island were dissected and prepared for scanning electron microscopy. The three molecular markers COI, 16S, and H3 were sequenced and analysed through maximum likelihood and Bayesian methods. The phylogenetic analyses and species delimitation tests clearly distinguished two species, *T. challengeriana* and *T. dantarti*, being widely-spread in the Southern Ocean, and endemic to Bouvet Island, respectively. Coloration seemed to be an unreliable character to differentiate among species since molecular data revealed both species can either have orange or white colour-morphotypes. This variability could be explained by pigment sequestration from the soft coral species they feed on. Morphological analyses reveal differences between Antarctic and Magellanic specimens of *T. challengeriana*, thus, we suggest the resurrection of *T. antarctica* Martens & Pfeffer, 1886 to encompass exclusively the Antarctic species. To progress further, additional molecular data from Magellanic specimens are required to definitely resolve their taxonomy and systematics.

## INTRODUCTION

The organisms composing Antarctic benthic fauna tend to present long life cycles, slow growth rates due to slow metabolism, and direct development; and this is particularly true for molluscs (Peck et al. 2006; Moles et al. 2017). All these common characteristics seem to be the consequence of the peculiar characteristics of the Southern Ocean (SO), e.g. low temperatures, relative stability in the frequency of physical disturbance, and pronounced seasonality (Dayton et al. 1992; Chown et al. 2015; Riesgo et al. 2015) aided by the onset of the Antarctic Circumpolar Current (ACC), ca. 25 Mya (Beu et al. 1997). During the late Eocene glacial periods, shelf fauna was completely impoverished with some species migrating into shelters (i.e. polynyas) and deep-sea waters, these being one of the major shelters for eurybathic species during the Last Glacial Maximum (Thatje et al. 2005). Certain taxa were able to re-colonize shallow waters during interglacial periods or when iceberg scouring wrecked the benthic communities and left free space available (Thatje et al. 2005). The deeper shelf of the Antarctic continent and the periodic destruction of benthic habitat on the shelf were hypothesized as natural evolutionary drivers towards eurybathy (i.e. capacity of species of living at a wide depth range), a widely shared feature of the Antarctic benthic fauna (Thatje et al. 2005; Allcock and Strugnell 2012). Numerous taxa present circum-Antarctic distributions due to the action of the ACC, the main responsible for the connectivity between populations due to the clockwise dispersion of larvae and/or adults around the SO (Thatje 2012; Riesgo et al. 2015). On the other hand, the Polar Front acts as a North-South barrier for water exchange above 1000 m depth (Clarke et al. 2005). The idea of the SO being isolated by the Polar Front has been challenged during the last years, revealing species connectivity and genetic flow with the adjacent areas (e.g. South Africa and the Magellanic region; Griffiths 2010, Chown et al. 2015).

Gastropods are one of the major taxa represented in the SO, with numerous species still being discovered (e.g. Moles et al. 2018, 2019; Fassio 2019; Layton et al. 2019). In the SO, nudibranchs are currently represented by less than a hundred recognized species (Moles 2016; De Broyer et al. 2019), although this species richness could increase with the application of molecular techniques. Among nudibranchs, the Dendronotida gathers several taxa with unassigned or unstable phylogenetic position (Goodheart et al. 2015). One of these taxa is the family Tritoniidae, among which the genus *Tritonia* Cuvier, 1798 appears to be the most speciose (WoRMS Editorial Board 2018). Currently, there are 35 valid species belonging to the genus *Tritonia*, and only three of them are found in the SO, with Antarctic, Sub-Antarctic, and Magellanic distributions, namely *T. dantarti* Ballesteros & Avila, 2006, *T. vorax* (Odhner, 1926), and *T. challengeriana* Bergh, 1884, respectively. *Tritonia vorax* was firstly described from South Georgia as *Duvaucelia vorax* by Odhner in 1926 and then transferred into *Tritonia* by Marcus in 1958 (Wägele 1995; Schrödl 2009). *Tritonia dantarti* was described in 2006 from Bouvet Island (Ballesteros and Avila 2006). *Tritonia challengeriana*, instead, was described for the first time in 1884 by Bergh from the Magellan Strait (Bergh, 1884). Since then, the latter species has been found in South Georgia, the Falkland Islands, Tierra del Fuego, and in several Antarctic locations (Antarctic Peninsula, Ross Sea, Scotia Arc; Wägele, 1995; Schrödl, 2003). Since its first description, several nominal species have been synonymized. In Antarctica, *T. antarctica* Pfeffer in Martens & Pfeffer, 1886, was first described by Pfeffer (1886) from South Georgia, and later ascribed to *T. challengeriana* by Odhner (1926). Years later, Wägele (1995) differentiated between Magellanic specimens which were identified as *T. challengeriana* and specimens occurring south of the Antarctic convergence, regarded as *T. antarctica*. This was based on the presence of oral lips and the absence of mantle glands in *T. antarctica*. However, Schrödl (1996) mentioned that oral lips may also be present in *T. challengeriana* from the Chilean Patagonia. Mantle glands were found in histological sections of *T. antarctica* from South Georgia although in much lower numbers than in *T. challengeriana* from the Magellan area, and this led to synonymize again *T. antarctica* with *T. challengeriana* (Schrödl 2003). According to Schrödl (2003, 2009), there are also other described species that are no longer valid and are considered synonyms of *T. challengeriana*, i.e. *Microlophus poirieri* Rochebrune & Mabille, 1889, *T. poirieri* Odhner (1926), and *T. australis* (Berg, 1898). The specimens collected for these studies were often limited to a single individual and thus these identifications might be unreliable (Wägele, 1995; Schrödl, 2003, 2009; Shields et al. 2009). Furthermore, until now, no molecular data are available for any of these species when given the wide range of distribution that *T. challengeriana* seems to present, the implementation of molecular tools could prove helpful to solve this phylogenetic conundrum. Here, we aim to combine molecular techniques, used here for the first time in this species complex, with detailed morpho-anatomical analysis to shed light into the long-term discussed systematics and taxonomy of Antarctic *Tritonia* species.

## MATERIAL AND METHODS

### Sample collection

Specimens were collected by Agassiz trawl, bottom trawl, and Rauschert dredge at the Sub-Antarctic Bouvet Island and the eastern Weddell Sea in 1998 during the ANT XV/3 (Gutt and Arntz 1999) and in 2003–2004 during the ANT XXI/2 cruises (Brey 2005) of the R/V Polarstern (Alfred Wegener Institute, Bremerhaven, Germany) (Fig. 1). The specimens of *Tritonia* spp. were collected at depths ranging from 130 to 789 m at 17 different stations (Suppl. Material 1). Specimens were photographed on board and preserved in either Karnovsky, 70% ethanol, or 10% formalin in seawater for morpho-anatomical analyses, or frozen and later transferred to 96% ethanol, for molecular analyses.

**Fig. 1.**
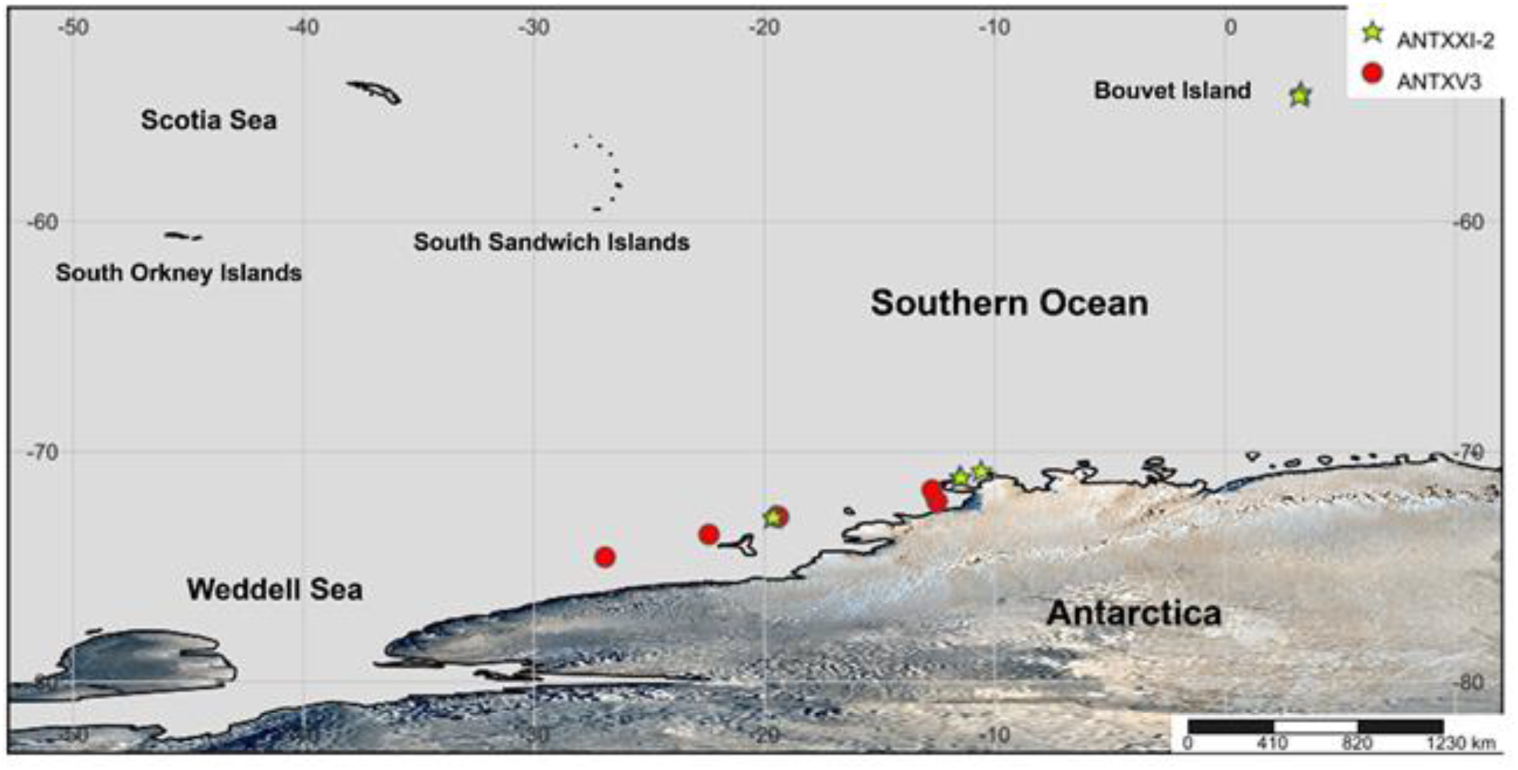
Map of the Western Antarctic region showing the sampling stations of the ANT XV/3 (red circles) and the ANT XXI/2 cruises (yellow stars). Source: http://www.simplemappr.net/#tabs=0

### DNA amplification and extraction

Total genomic DNA was extracted from foot tissue with the DNeasy Tissue Kit (Qiagen, Valencia, CA, USA) following the manufacturer’s protocol. Molecular markers included two fragments of the mitochondrial genes cytochrome *c* oxidase I (COI) and 16S *rRNA*, and the nuclear gene *histone-3* (H3). Partial sequences of the protein-encoding COI gene were amplified using the primers LCO1490 and HCO2198 (Folmer et al. 1994), the 16S gene was amplified using 16Sar-L and 16Sbr-H (Palumbi et al. 2002), and the H3 gene was amplified with H3AD5’3’ and H3BD5’3’ (Colgan et al. 1998). PCR amplifications were carried out in a 10 μL-reaction including 5.1 μL of Sigma dH_2_O, 3.3 μL REDExtract-N-Amp PCR ReadyMix (Sigma Aldrich, St. Louis, MO, USA), 0.3 μL of each primer, and 1 μL of genomic DNA, following standard protocols implemented in our lab (Moles et al. 2016). The PCR for COI consisted of an initial denaturation step at 95 °C for 3 min, 39 cycles of denaturation at 94 °C for 45 s, annealing at 48–50 °C for 30 s, extension at 72 °C for 2 min, and a final extending step at 72 °C for 10 min. The PCR program for 16S involved an initial denaturing step at 94 °C for 3 min, 39 cycles of denaturation at 94 °C for 30 s, annealing at 44–52 °C for 30 s, extension at 72 °C for 2 min, and a final extending step at 72 °C for 10 min. For H3 amplifications we used an initial denaturation step at 94 °C for 3 min, 35 amplification cycles (94 °C for 35 s, 50 °C for 1 min, and 72 °C for 1 min and 15 s), and a final extension at 72 °C for 2 min. Amplified products were sequenced at the UB Scientific and Technological Centers (CCiT-UB) on an ABI 3730XL DNA Analyzer (Applied Biosystems, CA).

### Phylogenetic analyses

Chromatograms were visualized, edited, and assembled in Geneious Pro 8.1.5 (Kearse et al., 2012). To check for contamination, sequences were compared against the GenBank database using the BLAST algorithm (Basic Local Alignment Search Tool; Altschul et al. 1990, http://www.ncbi.nlm.nih.gov). Single gene sequences were aligned with the MUSCLE algorithm and alignments were trimmed to a position at which more than 50% of the sequences had nucleotides. Missing positions at the ends were coded as missing data. We used GBlocks 0.91b on the final trimmed alignment for identifying and excluding blocks of ambiguous data in the single, non-codifying gene alignments of 16S, using both relaxed and stringent settings (Talavera and Castresana 2007).

The best-fit model of evolution (GTR + Г + I; Yang 1996) was chosen using the Akaike information criterion (AIC; Posada and Buckley 2004) implemented in jModelTest 2.1.7 (Posada 2008) with the selected partition for each gene.

For each gene a maximum-likelihood (ML) analysis was conducted, the final result was given by a concatenated alignment of all three genes. ML analyses were conducted using RAxML 8.1.2 (Stamatakis 2014), using a GTR model of sequence evolution with corrections for a discrete gamma distribution and invariable sites (GTR + Г + I; Yang 1996) was specified for each gene partition, and 500 independent searchers were conducted. Nodal support was estimated through bootstrap algorithm (500 replicates) using the GTR-CAT model (Stamatakis et al. 2008). The Bayesian inference (BI) was performed on the concatenated alignment of the three genes using MrBayes 3.2.5 (Ronquist et al. 2011). Two runs were conducted in MrBayes for 10 million generations, sampling every 2,000^th^ generation, using random starting trees. A 25% of the runs were discarded as burn-in after checking for stationarity with Tracer 1.7 (Rambaut et al. 2018). Bootstrap support (BS) and posterior probabilities (PP) were thereafter mapped onto the optimal tree from the independent searches. The tree was rooted using four selected Proctonotoidea species as sister group to the rest of the Dendronotoidea species included in this study (see Goodheart et al. 2015).

### Species delimitation tests

To examine the molecular distinctiveness of the different Antarctic *Tritonia* morpho-species we used ABGD via the web interface (at http://wwwabi.snv.jussieu.fr/public/abgd/abgdweb.html; accessed 23^rd^ September 2017). ABGD was run using the K80 calibrated index of genetic distance with transition/transversion ratio (TS/TV) equal to 2.0 and with a fasta file input of the COI alignment. We applied default values for *P*_min_, *P*_max_, and the relative gap (1.0). Additionally, with the same alignment, a GMYC (Fujisawa et al. 2013) analysis was performed. The ultrametric tree, necessary for the GMYC, was generated with BEAST (Suchard et al. 2018), with GTR G+I substitution model, and lognormal relaxed clock with rate 1.0, for 10 million generations. TreeAnnotator was used to discard 25% as burn-in. The GMYC was performed on the webserver (https://species.h-its.org/gmyc/) with single threshold parameters.

### Morphological analyses

Photographs of whole animals were taken with a Nikon d300 Sigma 105mm f 2.8–32. Total length (L) was measured aided by a calliper. Specimens were dissected sagittally with the aid of fine forceps under a stereomicroscope. Radula and jaws were obtained from the buccal bulb after dissolving the oral bulb’s soft tissue in a 10% NaOH solution for up to four hours and later rinsed with distilled water in ultrasound baths. The reproductive system was depicted, and the penial papilla extracted, and critical point dried prior to mounting on stubs with carbon sticky-tabs, as for the radulae and jaws, for scanning electron microscopy (SEM). The stubs were carbon-coated, and images were taken using a J-7100F Jeol scanning electron microscope at the UB Scientific and Technological Centers (CCiT-UB).

## RESULTS

The description of the 50 specimens collected allowed us to classify them into the two known species *Tritonia challengeriana* and *T. dantarti*, which have been studied in detail here for their morphology and anatomy, as well as for their phylogenetic relationships.

### Systematics

Class GASTROPODA Cuvier, 1795

Subclass HETEROBRANCHIA Burmeister, 1837

Order NUDIBRANCHIA Cuvier, 1817

Suborder CLADOBRANCHIA William & Morton, 1984

Family TRITONIIDAE Lamarck, 1809

Genus *Tritonia* Cuvier, 1798

Type species: *Tritonia hombergii* Cuvier, 1803

***Tritonia challengeriana* Berg, 1884**

(Figures 2A–G, 5A–C, 6A–F)

**Fig. 2.**
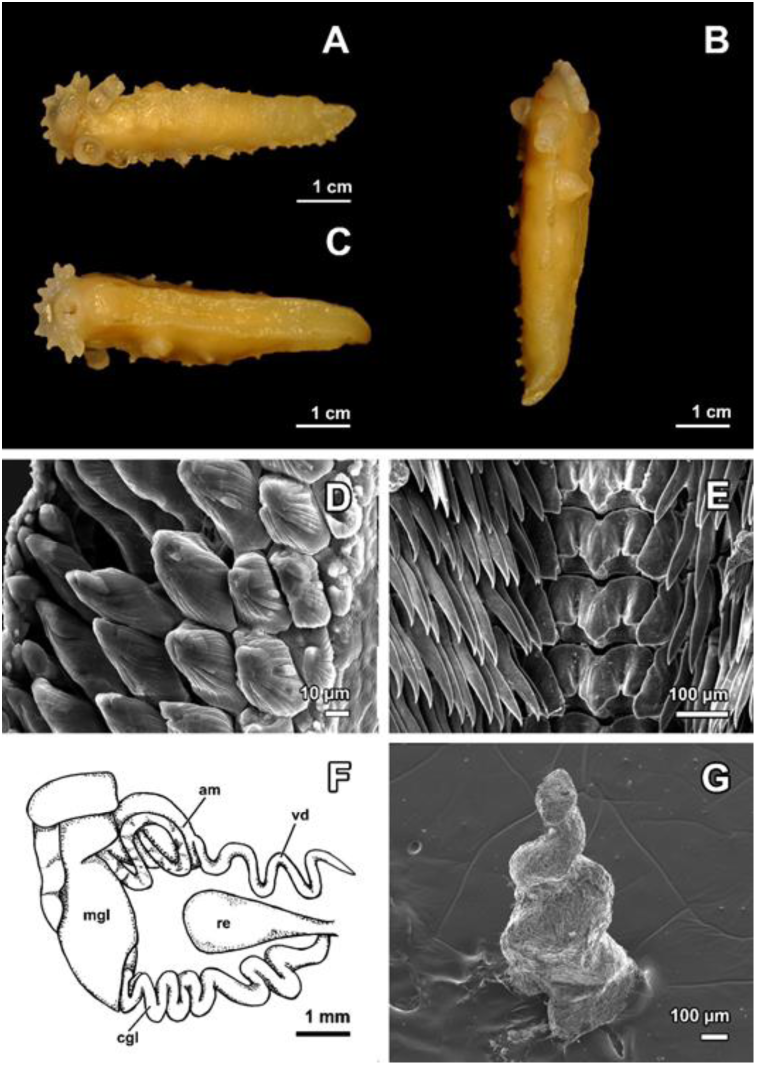
Preserved specimen of *Tritonia challengeriana* from the eastern Weddell Sea. (**A**) Dorsal view. (**B**) Lateral view. (**C**) Ventral view. (**D**) Detail of the jaw ornaments (SEM). (**E**) Scanning electron microscopy (SEM) of the radula showing the tricuspidated rachidian teeth, the first and subsequent lateral teeth. (**F**) Schematic drawing of the reproductive system. *am*, ampulla; *cgl*, capsule gland; *mgl*, mucous gland; *re*, seminal receptacle; *vd*, vas deferens (**G**) Detail of the penial papilla (SEM).

### Synonymy

*Tritonia challengeriana* Bergh, 1884: 45–47, pl. 11, figs. 16–19; Eliot 1907: 354–355; 1907: 3–5; Wägele 1995: 41–

45; Schrödl 2003: 97–101.

*Tritonia antarctica* Pfeffer in Martens & Pfeffer, 1886 112, pl. 3, figs. 6a,b; Wägele 1995: 21–46.

*Microlophus poirieri* Mabille & Rochebrune, 1889: 11–12, pl. 6, figs. 1a,b.

*Tritonia poirieri* (Mabille & Rochebrune, 1889): Wägele, 1995: 43;

*Candiella australis* Bergh, 1898

*Tritonia appendiculata* Eliot, 1905

*Tritonia australis* (Bergh): Dall 1909: 202.

*Duvaucelia challengeriana* (Bergh): Odhner, 1926: 35–37, pl. 1, fig. 14.

*Duvaucelia poirieri* (Rochebrune & Mabille): Odhner, 1926: 38–39

*Myrella poirieri* (Rochebrune & Mabille): Odhner, 1963: 51–52.

*Marionia cucullata* (Couthouy in Gould): Vicente & Arnaud, 1974: 539, figs. 6,7, pl. 3, figs. 1–3.

### Material examined

Out of the 37 specimens collected, 32 were frozen, two were fixed in 10 % formalin in seawater, two in 70 % ethanol, and one in 96 % ethanol. South of Vestkapp, eastern Weddell Sea, 73° 36.6’ S, 22°24.7’ W, 736 m depth: 1 spc., dissected and sequenced, T08, L = 25 mm, barcode MN651129. Halley Bay, 74°35.8’ S 26°55.0’ W, 789 m depth: 1 spc., dissected and sequenced, T10, L = 22 mm, barcode MN651131.North of Kapp Norvegia, 71°07.34’ S, 11°27.80’ W, 146 m depth: 1 spc., dissected, T25, L = 26 mm. Drescher Inlet, 72°05.18’ S, 19°38.62’ W, 598 m depth: 1 spc., dissected T28, L = 42 mm.

### External morphology (Fig. 2A–C)

Body length 22–42 mm after preservation (Table 1). Body wider dorsally than ventrally. Colour of preserved specimens milky white, beige-brownish when viscera seen by transparency; live specimens homogeneously white to orange. Dorsal mantle surface smooth, with subepithelial white knobs found mostly in posterior region of mantle. White pigmentation seen on notal margin, gills, and margin of rhinophoral sheath (Fig. 2A). Rhinophoral sheath broad; margin and plumes smooth. Gills ramified, dichotomous, large or small, from 6 to 19 per side, situated in parallel to each other. Oral veil prominent, bilobed or not. Five to ten velar processes present. Mouth surrounded by thick lips without distinct oral tentacles. Foot perimeter smaller than notal surface (Fig. 2C). Genital papilla found on right side of body. Anal opening placed at ½ of body length (Fig. 2B).

**Table 1.** Diagnostic characters of *Tritonia challengeriana, T. dantarti*, and *T. vorax* from this study and the literature. *n*.*r*. not reported.

|  | *T. challengeriana* | *T. challengeriana* | *T. dantarti* | *T. dantarti* | *T. vorax* |
| --- | --- | --- | --- | --- | --- |
| **Location** | Weddell Sea | Antarctic Peninsula,<br>Weddell Sea | Bouvet Island | Bouvet Island | South Georgia |
| **Body size after preservation (mm)** | 21–42 | 23–65 | 18–23 | 18–33 | 22–60 |
| **Colouration** | Beige-brownish,<br>milky white | White, yellow,<br>brown | Beige, milky white | Bright orange, white | White, pink, cream |
| **Dorsal mantle** | Smooth with<br>subepithelial knobs | Warty | Smooth with<br>subepithelial knobs | With crests | Smooth |
| **Oral lips** | Present | Present | Present | Present | Absent |
| **Velum** | Not prominent,<br>sometimes bilobed | Sometimes bilobed | Prominent or not,<br>bilobed | Wide, semitransparent,<br>bilobed | Bilobed |
| **Num. of velar processes** | 5–10 | <15 | 9–19 | <15 | >15 |
| **Num. of gill clusters per side** | 6–14 | 6–30 | 15–29 | 20–33 | 20–40 |
| **Size of gill clusters** | Large or small | Small or medium | Large or small | Large or small | Tiny or small |
| **Shape of gill clusters** | Ramified or<br>dichotomous | Digitated | Ramified, highly<br>digitated | Highly ramified | Dichotomous |
| **Salivary glands** | Thin, elongated | Long, ribbon-like | Large,<br>isodiametric | Small, elongated | Long, ribbon-like |
| **Rachidian tooth** | Broad, prominent monocuspidated | Broad, prominent central cusp | Broad, prominent monocuspidated | Broad, monocuspidated, central cusp and two lateral rounded protuberances | Broad, tricuspidated |
| **Radular formula** | 30–37 x (33–49).1.1.1.(33–49) | 31–46 x (37–63).1.1.1.(37–63) | 36–37 x (43–46).1.1.1.(43–46) | 30–38 x (35–48).1.1.1.(35–48) | 54–71 x (79–115).1.1.1.(79–115) |
| **Jaw/body proportions** | 0.21–0.3 | 0.25 | 0.25–0.28 | 0.27 | >0.4 |
| **Jaw ornaments** | Conical, striated, hooked | Conical, striated | Conical, striated, hooked | Conical, slightly curved | Conical, striated |
| **Penis** | Flagellated and/or conical | Digitiform to conical | Thin, flagellated and/or conical | n.r. | Digitiform to conical |
| **Gut contents** | Alcyonarian spicules, diatoms | *Cephalodiscus* | Alcyonarian spicules, diatoms, octocoral polyps | n.r. | Alcyonarian species, amphipods (*Gammaropsis*), tanaidaceans |
| **Reference** | This study | Wägele, 1995; Schrödl, 2003 | This study | Ballesteros and Avila, 2006 | Wägele, 1995 |

### Digestive system (Fig. 2D–E)

Oral lips smooth, large. Oral tube short. Oral bulb and pharynx thick. Jaws yellowish, curved towards inside; several rows of conic denticles, curved, striated, as border ornaments (Fig 2D). Jaw length ranging from 5 to 9 mm. Ratio jaw:body length ranging 0.21–0.3. Radular formula 30–37 x (33–49).1.1.1.(33–49); transparent in colour. Rachidian teeth presenting median prominent cusp, one smaller cusp found per each side (Fig 2E). Two different lateral teeth, with one short, broad cusp. Oesophagus running dorsally from pharynx. Salivary glands thin, elongated, running laterally from first half of body, then ventrally following oesophagus. Stomach situated ventrally. Intestine generally striated, originating dorsally from stomach, turning right, ending in anal opening.

### Reproductive system (Fig. 2F–G)

Reproductive system androdiaulic. Gonad large, wrinkled, covering digestive gland posteriorly. Gonoduct opening in ampulla connected by spermiduct to vas deferens. Bifurcation into vas deferens and oviduct not easily detected (Fig. 2F). Penial sheath terminal, conical, thin; penial papilla conical, slightly twisted (Fig. 2G). Seminal receptacle voluminous, with short duct; going shortly into genital opening. Granulated capsule gland at bottom of wide mucous gland, preceding short oviduct.

### Ecology

The specimens were collected from 146 to 789 m depth. Sclerites of alcyonarian octocorals were found in the gut contents of the three specimens studied (Fig.5 B–D).

### Distribution

Argentinian Patagonia (Marcus et al. 1969), Falkland Islands (Eliot 1907), Chilean Patagonia (Bergh 1884; Schrödl 1996) to Ancud Bay (Schrödl 1996), South Georgia (Odhner 1926), Adélie Land (Vicente and Arnaud 1994), Victoria Land (Ross Sea, Schiaparelli et al. 2006), eastern Weddell Sea (Wägele 1995; this study).

### Remarks

Most synonyms of genera and species to *T. challengeriana* were based on external morphological similarities (Wägele 1995; Schrödl 2003). For instance, Mabille and Rochebrune (1889) described *Microlophus poirieri* from Patagonia, Falkland Islands, and South Georgia, based only on their external morphology and colouration. This was later synonymized to *T. challengeriana* by Wägele (1995) based on their external morphology. *Marionia cucullata* was described from Adélie Land (Vicente and Arnaud, 1974). The similarity in oral lips’ shape and the low number of gills allowed Wägele (1995) to synonymize this genus and species to *T. antarctica. Tritonia appendiculata* Eliot, 1905 was described from Harbour of South Orkney, Scotia Bay, at 16 m depth. Its body colour was greenish-yellow, this is the only character that clearly differs from our specimens, since most of the morphological characters overlap with *T. challengeriana*. For instance, the species presents 19 gills per side and, on the dorsal surface, sub-epithelial knobs organized as “warts” were present (Eliot 1905). The oral veil presents twelve simple digitate processes and the lips are projected on each side of the mouth. The relation of jaw length (10 mm) to body length (51.5 mm) is 0.19 for *T. appendiculata*, while Wägele (1995) found a similar ratio for *T. challengeriana* 0.23, as in our study, thus our data support Wägele’s synonymy. *Tritonia poirieri* Mabille & Rochebrune, 1891, det. Odhner 1926, was found at Fitzroy Channel, at 14 m depth. The species body shape resembles that of *Doris*, with the notal margin bent downward. Other than the peculiar body shape, there were not enough differences to clearly identify *T. poirieri* as a distinct species from *T. challengeriana* (Wägele 1995).

Wägele (1995) differentiated Magellanic specimens of *T. challengeriana* from the specimens occurring south of the Polar Front, regarded as *T. antarctica* Martens & Pfeffer, 1886. The major difference was the presence of oral lips and mantle glands, exclusively found in *T. antarctica*. Later on, Schrödl (2003) described these two characters in *T. challengeriana* from Chilean Patagonia and synonymized it to *T. antarctica*. Our specimens are morphologically similar to the *T. antarctica* specimens described by Wägele (1995), with visible white knobs on the dorsal surface of the body and the presence of conspicuous oral lips. In fact, our description of *T. challengeriana* overlaps with the measurements and descriptions from Pfeffer (1886) and Wägele (1995), thus highlighting a major similarity to *T. antarctica* than to the Magellanic *T. challengeriana*. On the other hand, *Tritonia vorax* (Odhner, 1926) is found in South Georgia (Wägele 1995), Burdwood Bank, and the Chilean Patagonia (Odhner 1926; Schrödl 1996). Living specimens present a whitish to brownish colouration, with white or opaque white reticulations on the notal surface. Preserved specimens can be whitish, yellowish or pinkish and their notum can be more or less smooth. This species differs from *T. challengeriana* by having less number of gills, extremely large and strong jaws, which cause an elevated mediodorsal protuberance in between the rhinophores, and the lack of oral lips, with a higher jaws:body length ratio than *T. challengeriana* (Table 1). Differences between *T. challengeriana* and *T. dantarti* are discussed in the Remarks section below.

***Tritonia dantarti* Ballesteros & Avila, 2006**

(Figures 3A–G, 5D–E, 7A–B)

**Fig. 3.**
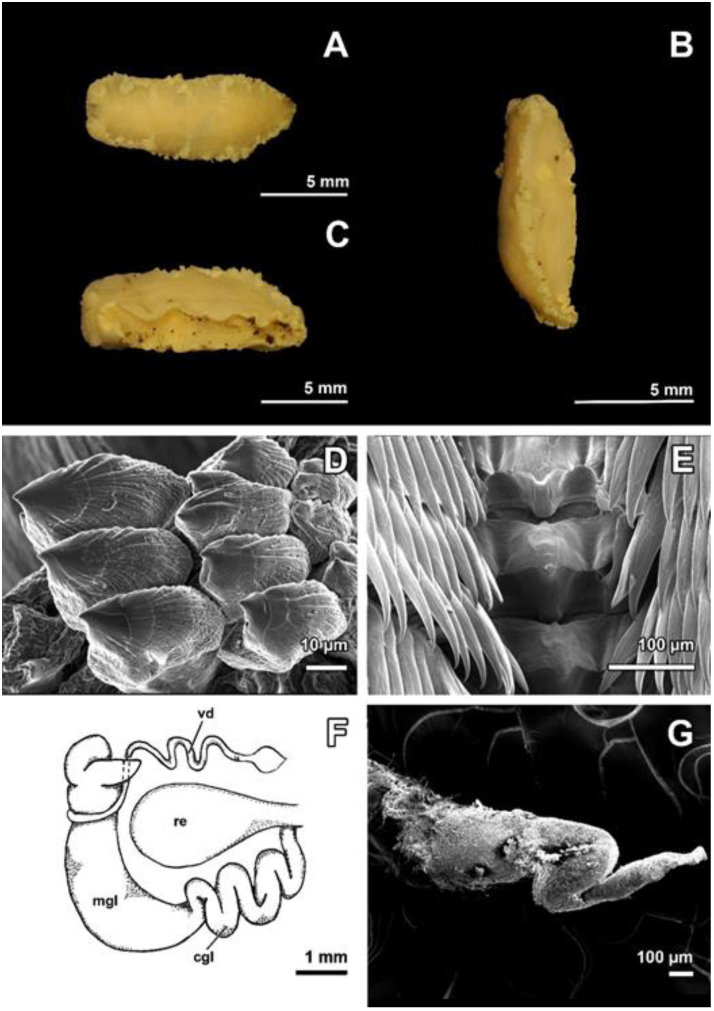
Preserved specimen of *Tritonia dantarti* from Bouvet Island. (**A**) Dorsal view. (**B**) Lateral view. (**C**) Ventral view. (**D**) Detail of the jaw ornaments (SEM). (**E**) Scanning electron microscopy (SEM) of the radula showing the rachidian teeth, the first and subsequent lateral teeth. (**F**) Schematic drawing of the reproductive system. *am*, ampulla; *cgl*, capsule gland; *mgl*, mucous gland; *re*, seminal receptacle; *vd*, vas deferens (**G**) Detail of the penial papilla (SEM)

### Material examined

Thirteen specimens collected at stations PS65/028-1 and PS65/029-1 in Bouvet Island. Six specimens were fixed in 70% ethanol, four were frozen, one in 96% ethanol, and two in Karnowsky. Bouvet Island, 54° 30.1’ S, 3°13.97’ W, 260 m depth: 1 spc, dissected and sequenced, T14.3, L = 18 mm, barcode MN651134; 54° 22.49’ S, 3°17.58’ W, 130 m depth: 1 spc, dissected, T16, L = 23 mm; 54° 22.49’ S, 3°17.58’ W, 130 m depth: 1 spc, dissected, T18.1, L = 20 mm, barcode.

### External morphology (Fig. 3A–C)

Body short, thick; 18–23 mm length. Colour beige to milky white in preserved specimens, living specimens sometimes completely white (Fig. 8A) or bright orange on dorsal surface, with warts forming a reticulation; white laterally (Fig. 8B). Dorsal mantle surface smooth with subepithelial white knobs. Notal margin unpigmented. Rhinophores with large sheath; smooth margin with emerging plumes. Single small gills or largely ramified, from 15 to 29 per side (Fig. 3A). Oral veil not prominent, bilobed or not; velar processes, short, nine to 19 in number. Lips thick, surrounding buccal bulb, without recognizable tentacles. Foot narrower than notum (Fig. 3C). Genital papilla on right side at 1/3 of body length. Anal opening at ½ of body length (Fig. 3B). Length and morphometrical data reported in Table 1.

### Digestive system (Fig. 3D–E)

Oral lips thick, smooth. Pharynx large, compact; hosting a pair of curved jaws, with a yellowish margin. Jaw denticles broad, conical, striated, hooked on top, arranged in several rows (Fig 3D). Jaw length ranging 5–6.5 mm. Jaw:body length ratio ranging from 0.25 to 0.28. Rachidian teeth broad, monocuspidated (Fig. 3E). Radular formula: 36–37 x (43–46)1.1.1.(43–46). Oesophagus running dorsally from pharynx. Salivary glands large, isodiametric, running laterally in first body half, then ventrally under oesophagus. Stomach situated ventrally. Intestine generally striated, originating dorsally from stomach, turning to right side, ending at anal opening.

### Reproductive system (Fig 3F–G)

Reproductive system situated between buccal bulb and digestive gland. Gonad brownish, warty, covering digestive gland. Genital papilla opening in ampulla, spermiduct could not be observed (Fig. 3F). Seminal receptacle wide. Penis thin, flagellated (Fig. 3G), occasionally conical. Penial papilla with conical shape. Mucus gland well developed, situated on top of entire system; granulated capsule gland preceding short oviduct, often convoluted.

### Ecology

Specimens of *T. dantarti* were collected on Bouvet Island at 130–134 m depth, in sea bottoms dominated by ophiuroids (e.g., *Ophionotus victoriae*), sea stars (*Porania antarctica*), holothuroids, sedentary polychaetes, hydroids, alcyonarians, different actinian species, amphipods, and pycnogonids. Gut contents showed that *T. dantarti* feeds on alcyonarians of the genus *Alcyonium* (Fig. 5A, 5E).

### Distribution

Northwest and southeast of Bouvet Island.

### Remarks

*Tritonia dantarti* is clearly distinguished from its counterpart *T. vorax* by the possession of oral lips, completely lacking in *T. vorax*. In *T. dantarti* the oral veil can be bilobed or not, while it is always bilobed in *T. vorax*. Moreover, *T. dantarti* presents lesser teeth rows and a monocuspidated rachidian tooth, while *T. vorax* presents a higher number of rows with a tricuspidated rachidian tooth. Additionally, the jaws:body length ratio is higher in *T. vorax* (Table 1).

*Tritonia dantarti* was described by possessing a conspicuous orange colouration in the dorsum of living specimens (see Fig. 6a,c,e in Ballesteros and Avila 2006; Fig. 7B in this study). This was, in fact, the main difference from *T. challengeriana*, but here molecular evidence of both white and orange colour-morphs is given for *T. dantarti* (see below). An additional diagnostic character is the presence of a warty reticulation in the notal surface of living specimens of *T. dantarti*, which has not been obviously observed in our preserved specimens, and is completely missing in *T. challengeriana*. Moreover, *T. challengeriana* generally presents fewer velar processes and fewer clusters of gills, but some overlap exists for both species, and a broad range of morphological differences are especially misleading in preserved specimens of both species. Our results agree with previous descriptions for both species (Wägele, 1995; Schrödl, 2003; Ballesteros and Avila, 2006).

### Phylogenetic analyses

The total dataset contained 41 specimens of *Tritonia* and 17 closely related outgroup taxa (Suppl. Material 2). The concatenated alignment consisted of 1,415 characters, including COI with 3^rd^ codon position (ca. 601 bp), 16S unmodified (ca. 486 bp), and H3 with 3^rd^ codon position (ca. 328 bp). The best-fit evolutionary models and parameters were calculated by jModeltest and Gblocks (Suppl. Material 3).

ML and BI analyses recovered a tree with maximum support for both *T. challengeriana* specimens from the Weddell Sea and the only sequenced specimen from the Ross Sea (PP = 1, BS = 100), and for *T. dantarti* including only the Sub-Antarctic specimens from Bouvet Island (PP = 1, BS = 98; Fig. 4). The GenBank specimen labelled as *T. antarctica* (voucher number CASIZ171177) clusters here with our specimens of *T. dantarti*, and thus might be considered a missidentification. Sister to both SO species sequenced we found the North Pacific *T. festiva*. The SO species clustered in a clade with highly supported clusters of different *Tritonia* species. We recovered the unidentified *Tritonia* sp. 3, *Tritonia* sp. 6, *Tritonia* sp. 7, and *Tritonia* sp. G all in a well-supported clade with all sequenced *Marionia* species. The relationships of the Antarctic monotypic *Tritoniella belli* were not clearly found in our analyses. The relationship among *Bornella, Marionia, Tritoniella*, and *Tritonia* clades was not recovered in this study.

**Fig. 4.**
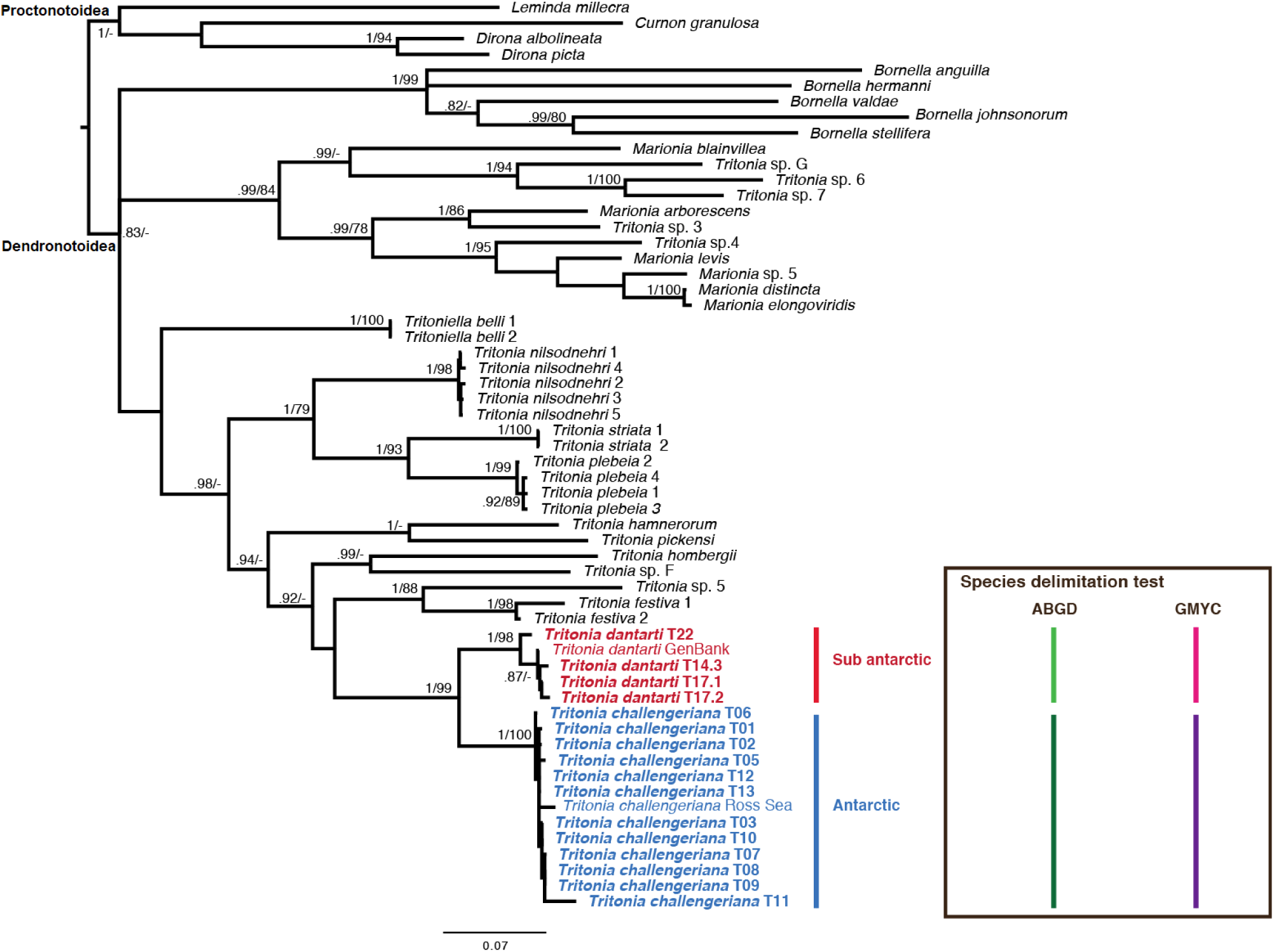
Phylogenetic tree of *Tritonia* species and outgroup species considered using Bayesian inference (BI) and maximum likelihood (ML) on the combined COI, 16S, and H3 datasets. Numbers on the nodes indicate posterior probability values (BI) and bootstrap support values (ML). The sequences generated in our lab are depicted in bold. The *T. dantarti* GenBank specimen placed in the red cluster is registered in GenBank as *T. antarctica*. (Voucher n. CASIZ171177). In the box, the results of the ABGD (green) and GMYC (purple) analyses are represented as bars, distinguishing the two SO species groups studied.

The ABGD analyses additionally supported the taxonomic classification of *T. challengeriana* and *T. dantarti* with an intraspecific variation of 1.7 and 1.9 % on average, respectively; whereas their interspecific variation ranged from 12 to 14 %. Intraspecific variation within other *Tritonia* species considered in this study range from 0 to 7 %, while their interspecific variations ranges approx. 9.1–25.7 % (Table 2). We have chosen to not consider in this species delimitation tests, the unidentified *Tritonia* spp. (Sup. Material 2) due to a possible misinterpretation of the specimens, that may belong to the genus *Marionia* (Fig. 4). The GMYC analysis also recovers two distinct species groups belonging to *T. challengeriana* and *T. dantarti*, in accordance to the ABGD and the phylogenetic tree (Suppl. Material 4).

**Table 2.** Kimura two-parameter distances (K2P) between and within groups for the putative species of *Tritonia* included in our analyses; *n*.*c*. non-computable since there was a single sequence available.

| Between groups |  |  |  |  |  |  |  |  |  | Within groups |  |
| --- | --- | --- | --- | --- | --- | --- | --- | --- | --- | --- | --- |
|  | 1 | 2 | 3 | 4 | 5 | 6 | 7 | 8 | 9 |  |  |
| 1 *T. challengeriana* |  |  |  |  |  |  |  |  |  | 1 | 0–1.5 |
| 2 *T. dantarti* | 7.9–14 |  |  |  |  |  |  |  |  | 2 | 0–0.01.9 |
| 3 *T. festiva* | 12.6–20.5 | 17.7–19 |  |  |  |  |  |  |  | 3 | 0–0.13.4 |
| 4 *T. hamnerorum* | 11.1–18.1 | 18.4–20 | 21.1–21.3 |  |  |  |  |  |  | 4 | n.c |
| 5 *T. hombergii* | 12.5–22.3 | 20.5–22.5 | 18.7–19.1 | 22.7 |  |  |  |  |  | 5 | n.c |
| 6 *T. nilsodnheri* | 8.9–22.5 | 11.4–21.6 | 11.1–21 | 9.2–16.8 | 10– 21.3 |  |  |  |  | 6 | 0–2 |
| 7 *T. pickensi* | 19.4–25.7 | 22.6–24.1 | 21.1–21.3 | 17.2 | 21.7 | 10.1–20.5 |  |  |  | 7 | n.c. |
| 8 *T. plebeia* | 12.4–22.6 | 21.7–23.3 | 21.4–22.3 | 20.2–20.6 | 23.5–23.9 | 9.1–18.9 | 19.9–20.8 |  |  | 8 | 0.1–7 |
| 9 *T. striata* | 13.4–23.6 | 21.2–23 | 20.1–20.3 | 21.3 | 24 | 10–20.6 | 21.9 | 15.6–21.9 | 16.6 | 9 | 0.00 |

## DISCUSSION

### Taxonomy and morphology of Antarctic *Tritonia* species

The specimens analysed in this study from the high Antarctic belonged to the only current valid species *Tritonia challengeriana*, while the specimens from Bouvet Island belonged to *T. dantarti*. Phylogenetic analyses and species delimitation tests recovered these two species with a strong support (Fig. 4), including the specimens of *T. challengeriana* from the Weddell Sea and the only sequenced specimen from the Ross Sea (PP = 1, BS = 100), and the specimens of *T. dantarti* from Bouvet Island (PP = 1, BS = 98). Morpho-anatomical analyses showed that, on the dorsal body surface in living specimens of *T. dantarti* warts and reticulation are visible. Nonetheless, the bright orange colouration (Ballesteros and Avila 2006) may no longer be a valid diagnostic character, since both milky-white and orange colour-morphotypes from Bouvet Island were found here, as it has been described for *T. challengeriana* from both South America and high Antarctic regions (Figs. 5–6). These results were supported for our molecular analyses. Besides this, no other clear diagnostic characters were found in the morpho-anatomical analyses to allow the discrimination among these two species. For instance, shape and body measurements, the number of velar processes, the shape and number of gills, the radular formula, and the shape of the jaws are not quite discernible between *T. dantarti* and *T. challengeriana*. In fact, both species overlap in the range of the aforementioned characters (Table 1), as also reported by Wägele (1995), Schrödl (2003), and Ballesteros and Avila (2006). Nonetheless, *T. challengeriana* seems to present lesser oral tentacles and gill clusters than *T. dantarti*, but still this might be subjected to ontogenetic development

The validity of *T. antarctica* has been questioned in a few studies (see remarks section of *T. challengeriana*). Wägele (1995) sustained the existence of *T. antarctica* for the presence, in Antarctic specimens, of subepithelial glands (externally visualized as knobs), which were lacking in Magellanic specimens. Later on, Schrödl (2003) suggested the contrary, showing that the glands were present on the dorsal surface of the specimens from the Magellanic area, even if sporadically and in a lower number. Our specimens seem to be similar to the *T. antarctica* described by Wägele (1995). Pictures of living specimens from the Magellanic region (Fig. 6A–C) do not show visible knobs, which are easily detectable on specimens from Antarctica (Fig. 6D–F). Unfortunately, we cannot confirm the validity of *T. antarctica*, since there are no molecular data available for *T. challengeriana* from the Magellanic region to date. Southern American material and additional samples from around Antarctica could be very useful to shed light into the Southern Hemisphere *Tritonia* species systematics. However, the morphological analysis suggests that *T. challengeriana* and *T. antarctica* could be considered to be different species, given the evidence of the visible knobs on the dorsal surface present on Antarctic specimens.

**Fig. 5.**
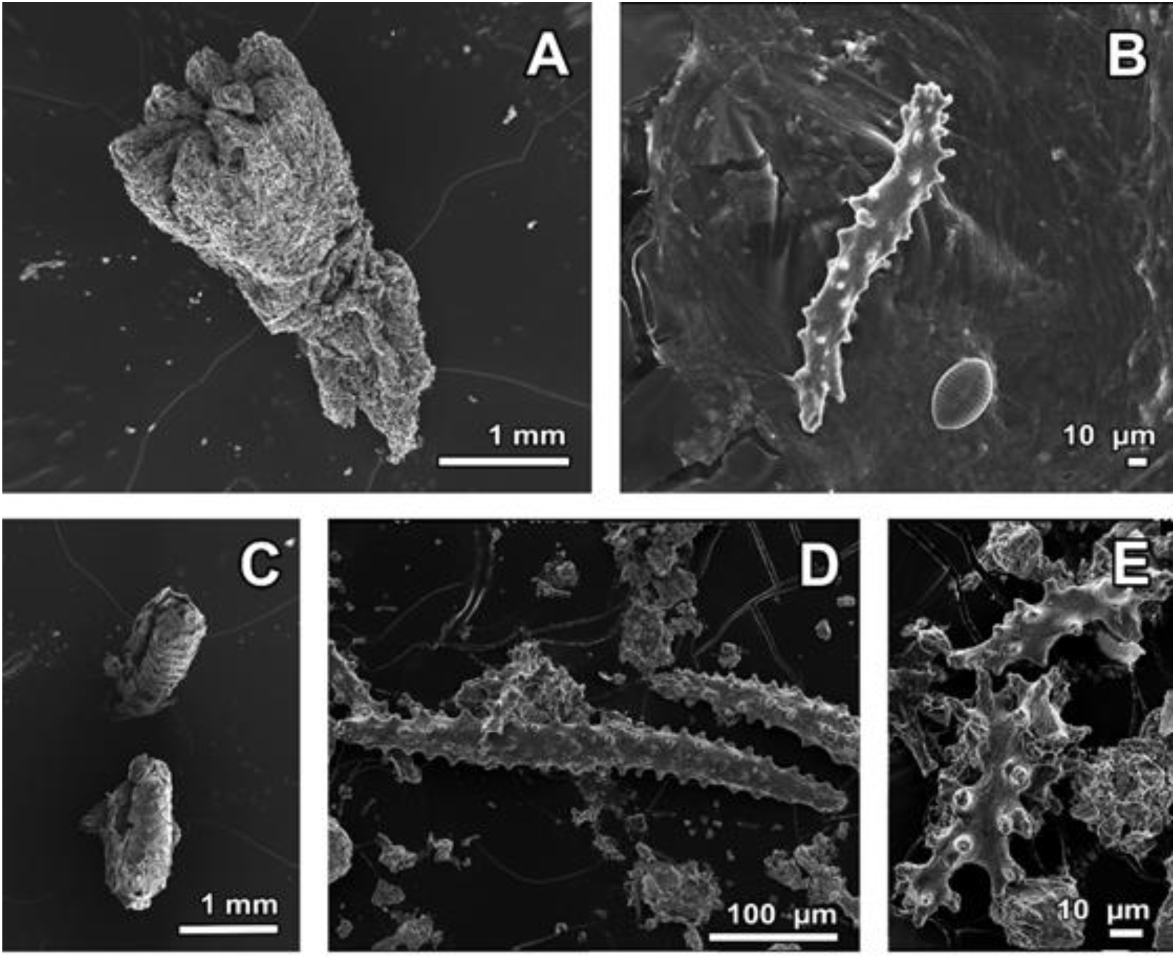
Gut content found in the intestine of the examined specimens. (**A–C**) Octocoral structures found in *Tritonia challengeriana*. (**B**) Alcyonarian sclerites and diatom found in *T. challengeriana*. (**D–E**) Alcyonarian spicules found in *T. dantarti*.

**Fig. 6.**
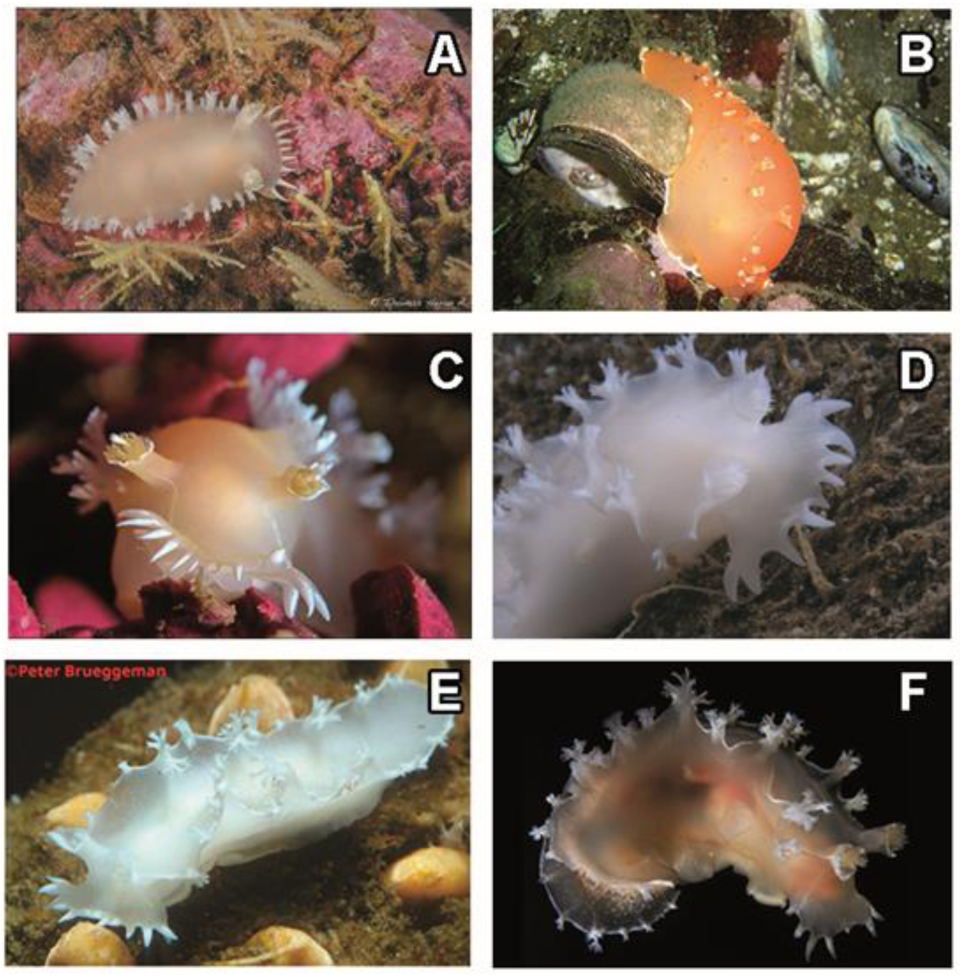
Underwater photographs of *Tritonia challengeriana* from its current range of distribution. (**A**) Puerto Raúl Marín Balmaceda, Chile (photograph by T. Heran). (**B**) Comau Fjord, Chile (photograph by D. Thompson). (**C**) Punta Porra, Chile (photograph by T. Heran). (**D**) Ross Sea, Antarctica (photograph by S. Harper) (**E**) Ross Sea, Antarctica (photograph by P. Brueggeman). (**F**) Antarctic Peninsula, Antarctica (photograph by G. Giribet).

**Fig. 7.**
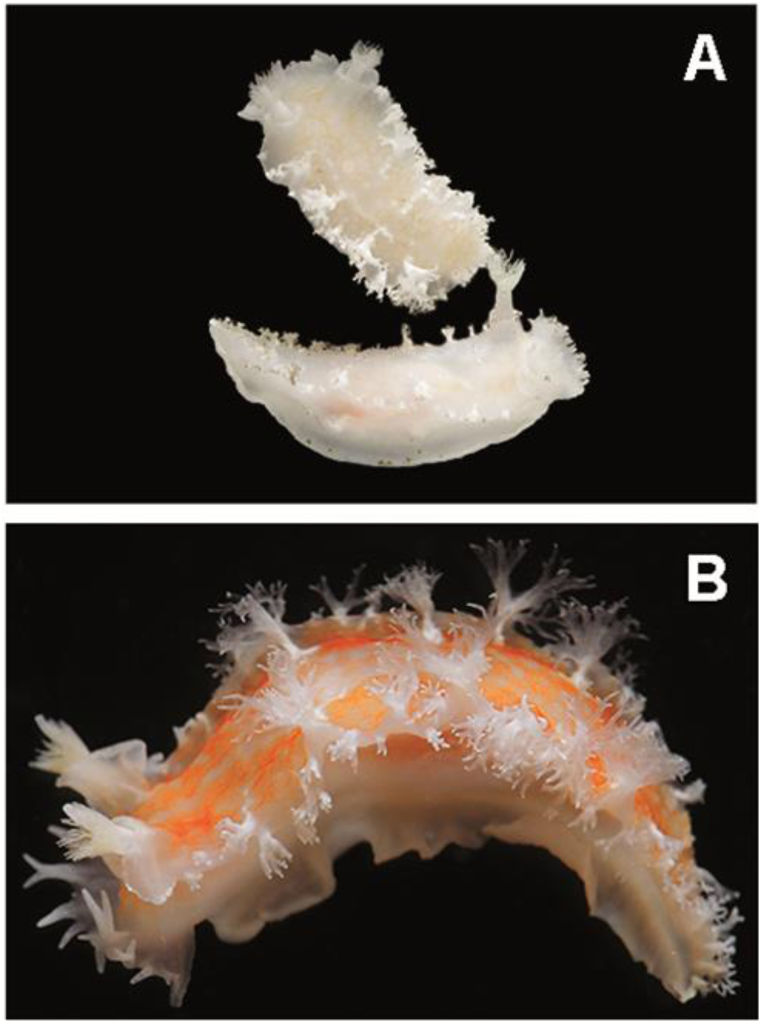
*Tritonia dantarti* specimens from Bouvet Island (photographs by M. Ballesteros). (**A**) Two specimens collected a 260 m of depth from Bouvet Island displaying whitish colouration (T14.3 and T14.4). (**B**) Specimen from Bouvet Island with orange colouration, collected at 130 m of depth (T15.1).

### The colouration issue

Members of the family Tritoniidae feed almost exclusively on octocorals, including sea pens, alcyonarian soft corals, and gorgonians, sometimes being cryptic in shape and colouration upon them (García-Matucheski and Muniain 2011). In the SO, *Tritonia* species feed mostly on alcyonarian soft corals (Schrödl 2003; Wägele 1995; García-Matucheski and Muniain 2011). Here, we found soft-coral sclerites in the gut contents of both *T. challengeriana* and *T. dantarti*. The *Alcyonium* species living in the SO are *A. antarcticum* Wright & Studer, 1889, *A. grandis* Casas, Ramil & van Ofwegen, 1997, *A. haddoni* Wright & Studer, 1889, *A. paucilobatum* Casas, Ramil & van Ofwegen, 1997, *A. sollasi* Wright & Studer, 1889, and *A. southgeorgensis* Casas, Ramil & van Ofwegen, 1997, and they can all present a yellow, cream or orange colouration, while they tend to be brighter in the Magellanic region (Casas et al. 1997). Through evolution, and related to the loss of the shell, nudibranchs have developed a plethora of defensive strategies against predators (Avila et al. 2018). These defences include chemicals (natural products), which can be either *de novo* synthesized by the own slug or gathered from their prey (*i*.*e*. kleptochemistry). An example of kleptochemistry in Antarctica is found in *Tritoniella belli* Eliot, 1907 which obtains its defensive natural products from its prey, the anthozoan *Clavularia frankliniana* Roule, 1902 (McClintock et al. 1994). Some dietary metabolites can be brightly coloured pigments, as for some *Alcyonium* spp. natural products (Abdel-Lateff et al. 2019), and these may provide an additional mimetic defensive strategy on top of the chemical deterrence of the slug. Although the development of a bright colouration may sometimes represent a warning mechanism (i.e. aposematism; Aguado and Marin 2007; Haber et al. 2010; Avila et al. 2018), the bright orange colouration found in both *T. challengeriana* and in *T. dantarti* may not represent an aposematism mechanism, since the majority of visually guided predators, such as fishes or decapods, are not especially diversified in Antarctica (De Broyer et al. 2011; discussed in Moles 2016). Nevertheless, some evidence is given for the defensive nature against sympatric sea star predators of *T. challengeriana*, although the compounds have not been identified yet (Avila et al. 2018). This strategy has been proved in other Antarctic species, such as *Bathydoris hodgsoni* Eliot, 1907 and *T. belli* (McClintock et al. 1994; Avila et al. 2000). We propose here that the colouration in *T. challengeriana* and *T. dantarti* varies locally, in direct relation to diet, and therefore cannot be used as a diagnostic character for these species.

### Distribution and cryptic speciation

From our molecular analyses, while *T. dantarti* seems to have a restricted, endemic distribution in Bouvet Island, *T. challengeriana* seems to present a disjunct (i.e. found in both the Weddell and Ross seas) and probably circumpolar distribution. This distribution could be partially explained by the action of the ACC (Thatje 2012). When the Drake Passage was narrower, the ACC flow was particularly intense in the Antarctic region, carrying adults or egg masses attached to floating debris (i.e. rafting phenomenon) to new habitats all around Antarctica (i.e. circumpolar distribution), where they could, through genetic drift and selection (Allcock and Strugnell 2012), diverge sufficiently to either yield a new species by allopatric speciation – which might have been the case for the restricted *T. dantarti* – or widening its geographical range, as for *T. challengeriana*. Even if the rafting phenomenon allowed a long-distance dispersal in organisms that not produce free-swimming larvae (Thatje 2012) during glacial cycles, the existence of polynyas, i.e. shelters and regions where the ice shelf did not cover homogeneously the shelf, acted as refugia (Thatje et al. 2005; Fraser et al. 2014; Chown et al. 2015) and species may have gone through a process of isolation, which led to cryptic speciation (Wilson et al. 2009). In fact, because of the existence of cryptic species, the current species richness of gastropods in the Antarctic and Sub-Antarctic regions is higher than previously thought (Linse et al. 2007). Likewise, new cephalaspidean molluscs with low character displacement have been recently described based on molecular data in the same region (Moles et al. 2017, 2019). The nudibranch *D. kerguelenensis* seems to also present this trend; molecular data evidenced a complex genetic structure that suggests much diversity than a single recognised species (Wilson et al. 2009). This hypothesis is corroborated by the wide variety of natural products used against predators, but due to the lack of morphological analyses the taxonomy of *D. kerguelenensis* is still not solved (Wilson et. al. 2013). Both *T. challengeriana* and *T. dantarti* are clearly two different species, but the relationship between *T. challengeriana* specimens from the Antarctic and the Magellanic regions remains still unclear, thus the validity of the *T. antarctica* requires further systematic work. Additional samples from other locations in Antarctica, Sub-Antarctic Islands, and South America are urgently needed to shed light on the systematics of the group. Sampling in poorly known areas of the SO, such as the Amundsen Sea or the western Weddell Sea, and the continental shelves underneath floating ice shelves (Griffiths 2010), with the increasing application of molecular techniques and complementary molecular markers with higher resolution (e.g. EPIC markers, microsatellites, and/or genome-or transcriptome-derived SNPs; Riesgo et al. 2015; Moles et al. 2019) are required to further evaluate cryptic speciation and increasing our knowledge on the biodiversity of most invertebrate taxa in the SO.

## ACKNOWLEDGMENTS

We are indebted to Prof W. Arntz, T. Brey, and the crew of R/V Polarstern, for allowing the participation of C. Avila in the Antarctic cruises ANT XV/3 and ANT XXI/2 (AWI, Bremerhaven, Germany). E. Prats (CCiT-UB, Barcelona, Spain) is acknowledged for her help during the SEM sessions. We also thank M. Ballesteros, P. Brueggeman, G. Giribet, S. Harper, T. Heran, and D. Thompson for kindly providing the pictures of the live animals. Funding was provided by the Spanish government through the ECOQUIM (REN2003-00545, REN2002-12006-E ANT) and DISTANTCOM (CTM2013-42667/ANT) Projects. M.E. Rossi was supported by an Erasmus + grant from La Sapienza, University of Rome, at the University of Barcelona. This paper is part of the AntEco (State of the Antarctic Ecosystem) Scientific Research Programme.

## REFERENCES

Aguado, F. and Marin, A. (2007). Warning coloration associated with nematocyst-based defences in aeolidiodean nudibranchs. Journal of Molluscan Studies, 73(1), 23–28.

Allcock, A.L. and Strugnell, J.M. (2012). Southern Ocean diversity: New paradigms from molecular ecology. Trends in Ecology and Evolution, 27(9), 520–528.

Altschul, S.F., Gish, W., Miller, W., Myers, E.W. and Lipman, D.J. (1990). Basic local alignment search tool. Journal of Molecular Biology, 215, 403–410.

Arntz, W. and Gutt, J., (1999). The expedition ANTARKTIS XV/3 (EASIZ II) of RV” Polarstern” in 1998. Berichte zur Polarforschung (Reports on Polar Research), 301.

Arntz, W. and Brey, T. (2005). The expedition ANTARKTIS XXI/2 (BENDEX) of RV” Polarstern” in 2003/2004. Berichte zur Polar-und Meeresforschung (Reports on Polar and Marine Research), 503.

Arntz, W.E., Thatje, S., Linse, K., Avila, C., Ballesteros, M., Barnes, D.K., Cope, T., Cristobo, F.J., De Broyer, C., Gutt, J. and Isla, E. (2006). Missing link in the Southern Ocean: sampling the marine benthic fauna of remote Bouvet Island. Polar Biology, 29 (2), 83–96.

Avila, C., Iken, K., Fontana, A. and Cimino, G. (2000). Chemical ecology of the Antarctic nudibranch *Bathydoris hodgsoni* Eliot, 1907: defensive role and origin of its natural products. Journal of Experimental Marine Biology and Ecology, 252(1), 27–44.

Avila, C., Taboada, S. and Núñez-Pons, L. (2008). Antarctic marine chemical ecology: what is next?. Marine Ecology, 29(1), 1–71.

Avila, C., Núñez-Pons, L. and Moles, J. (2018). From the Tropics to the Poles: chemical defense strategies in sea slugs (Mollusca: Heterobranchia). In: Puglisi MP, Becerro MA, editors. Chemical ecology: the ecological impacts of marine natural products (pp. 71–163). Boca Raton, CRC Press.

Ballesteros, M. and Avila, C. (2006). A new tritoniid species (Mollusca: Opisthobranchia) from Bouvet Island. Polar Biology, 29(2), 128–136.

Beu, A G., Griffin, M. and Maxwell, P. A. (1997). Opening of Drake Passage gateway and Late Miocene to Pleistocene cooling reflected in Southern Ocean molluscan dispersal: evidence from New Zealand and Argentina. Tectonophysics, 281(1–2), 83–97.

Casas, C., Ramil, F. and van Ofwegen, L.P. (1997). Octocorallia (Cnidaria: Anthozoa) from the Scotia Arc, South Atlantic Ocean: The genus *Alcyonium* Linnaeus,1758. Zoologische Mededelingen, 71(26), 299–311.

Chown, S.L., Clarke, A., Fraser, C.I., Cary, S.C., Moon, K.L. and McGeoch, M.A. (2015). The changing form of Antarctic biodiversity. Nature, 522(7557), 431.

Dall, W.H. (1909). Report on a collection of shells from Peru, with a summary of the littoral marine Mollusca of the Peruvian Zoological Province. Proceedings of the U. S. National Museum 37, 147–294.

Dayton PK, Mordida BJ, Bacon F (1994) Polar marine communities. American Zoologist 34, 90–99.

De Broyer, C. and Danis, B., 2011. How many species in the Southern Ocean? Towards a dynamic inventory of the Antarctic marine species. Deep sea research Part II: Topical studies in oceanography, 58(1-2), pp.5–17.

De Broyer, C., Clarke, A., Koubbi, P., Pakhomov, E., Scott, F., Vanden Berghe, W. & Danis, B. 2019. The SCAR-MarBIN Register of Antarctic Marine Species (RAMS), [06/04/2016]. World Wide Web electronic publication. Available online at http://www.scarmarbin.be/scarramsabout.php.

Eliot, C. 1905. The Nudibranchiata of the Scottish National Antarctic Expedition. Transactions of the Royal Society of Edinburgh, 41, 519–532.

Eliot, C. (1907). Nudibranchs from New Zealand and the Falkland Islands. Proceedings of the Malacological Society of London, 7, 350–361.

Fassio, G., Modica, M.V., Alvaro, M.C., Buge, B., Salvi, D., Oliverio, M. and Schiaparelli, S., (2019). An Antarctic flock under the Thorson’s rule: Diversity and larval development of Antarctic Velutinidae (Mollusca: Gastropoda). Molecular Phylogenetics and Evolution, 132, 1–13.

Fraser, C.I., Terauds, A., Smellie, J., Convey, P. and Chown, S.L. (2014). Geothermal activity helps life survive glacial cycles. Proceedings of the National Academy of Sciences, 111(15), 5634–5639.

García-Matucheski, S. and Muniain, C. (2011). Predation by the nudibranch *Tritonia odhneri* (Opisthobranchia: Tritoniidae) on octocorals from the South Atlantic Ocean. Marine Biodiversity, 41(2), 287–297.

Goodheart, J.A., Bazinet, A.L., Collins, A.G., and Cummings, M.P. (2015). Relationships within Cladobranchia (Gastropoda: Nudibranchia) based on RNA-Seq data: an initial investigation. Royal Society Open Science, 2(9), 150196.

Griffiths, H. J. (2010). Antarctic marine biodiversity–what do we know about the distribution of life in the Southern Ocean?. PloS ONE, 5(8), e11683.

Griffiths, H., and Grant, S. A. (2014). Biogeographic atlas of the Southern Ocean. C. De Broyer, and P. Koubbi (Eds.). Cambridge: Scientific Committee on Antarctic Research.

Haber, M., Cerfeda, S., Carbone, M., Calado, G., Gaspar, H., Neves, R., Maharajan, V., Cimino, G., Gavagnin, M., Ghiselin, M.T. and Mollo, E. (2010). Coloration and defense in the nudibranch gastropod *Hypselodoris fontandraui*. The Biological Bulletin, 218(2), 181–18.

Jörger, K.M., Schrödl, M., Schwabe, E. and Würzberg, L. (2014). A glimpse into the deep of the Antarctic Polar Front– Diversity and abundance of abyssal molluscs. Deep Sea Research Part II: Topical Studies in Oceanography, 108, 93–100.

Jossart, Q., Moreau, C., Agüera, A., De Broyer, C. and Danis, B. (2015). The Register of Antarctic Marine Species (RAMS): a ten-year appraisal. ZooKeys, 2015(524), 137–145.

Kearse, M., Moir, R., Wilson, A., Stones-Havas, S., Cheung, M., Sturrock, S., Buxton, S., Cooper, A., Markowitz, S., Duran, C., Thierer, T., Ashton, B., Mentjies, P., and Drummond, A. (2012). Geneious Basic: an integrated and extendable desktop software platform for the organization and analysis of sequence data. Bioinformatics, 2828(12), 1647–1649.

Layton, K. K., Rouse, G. W., & Wilson, N. G. (2019). A newly discovered radiation of endoparasitic gastropods and their coevolution with asteroid hosts in Antarctica. BMC evolutionary biology, 19(1), 180.

Lin, C.P. and Danforth, B.N. 2004. How do insect nuclear and mitochondrial gene substitution patterns differ? Insights from Bayesian analyses of combined datasets. Molecular Phylogenetics and Evolution, 30(3), 686–702.

Linse, K., Cope, T., Lörz, A.N. and Sands, C. (2007). Is the Scotia Sea a centre of Antarctic marine diversification? Some evidence of cryptic speciation in the circum-Antarctic bivalve *Lissarca notorcadensis* (Arcoidea: Philobryidae). Polar Biology, 30(8), 1059–1068.

Marcus, E. (1959). Lainellariacea and Opisthohranchia. Reports of the Lund University Chile Expedition, 55(9), 1–133.

Marcus, E.D.B.R., Marcus, E. and Kirsteuer, E. (1969). Opisthobranchian and lamellarian gastropods collected by the” Vema”. American Museum Novitates, 2368.

McClintock, J.B., Baker, B.J., Slattery, M., Heine, J.N., Bryan, P.J., Yoshida, W., Davies-Coleman, M.T. and Faulkner, D.J. (1994). Chemical defense of common Antarctic shallow-water nudibranch *Tritoniella belli,* Eliot (Mollusca: Tritonidae) and its prey, *Clavularia frankliniana,* Rouel (Cnidaria: Octocorallia). Journal of Chemical Ecology, 20, 3361.

Moles, J. (2016). Antarctic heterobranch molluscs: diving into their challenging ecology, taxonomy, and systematics. Doctoral thesis, Universitat de Barcelona.

Moles, J., Wägele, H., Ballesteros, M., Pujals, Á., Uhl, G. and Avila, C. (2016). The end of the cold loneliness: 3D comparison between *Doto antarctica* and a new sympatric species of *Doto* (Heterobranchia: Nudibranchia). PLoS ONE, 11, e0157941.

Moles, J., Wägele, H., Cutignano, A., Fontana, A., Ballesteros, M. and Avila, C. (2017). Giant embryos and hatchlings of Antarctic nudibranchs (Mollusca: Gastropoda: Heterobranchia). Marine Biology, 164(5), 114.

Moles, J., Avila, C. and Malaquias, M. A. E. (2018). Systematic revision of the Antarctic gastropod family Newnesiidae (Heterobranchia: Cephalaspidea) with the description of a new genus and a new abyssal species. Zoological Journal of the Linnean Society, 183(4), 763–775.

Moles, J., Avila, C. and Malaquias, M. A. E. (2019). Unmasking Antarctic mollusc lineages: novel evidence from philinoid snails (Gastropoda: Cephalaspidea). Cladistics, 35(5), 487–513.

Odhner, N.H.J. 1926. The Opisthobranchien (Vol. 2, No. 1). PA Norstedt & Söner.

Odhner N.H.J. (1936). Nudibranchia, Dendronotacea; A revision of the system. Mémoires du Musée Royal l’Histoire Naturelle de Belgique, 2, 1056–1128.

Odhner, N.H.J. 1963. On the taxonomy of the family Tritoniidae (Mollusca: Opisthobranchia). Veliger, 6, 48–52.

Peck LS, Clarke A, Chapman AL (2006) Metabolism and development of pelagic larvae of Antarctic gastropods with mixed reproductive strategies. Marine Ecology Progress Series, 318, 213–220.

Posada, D. and Buckley, T.R. (2004). Model selection and model averaging in phylogenetics: advantages of Akaike information criterion and Bayesian approaches over likelihood ratio tests. Systematic Biology, 53(5), 793–808.

Posada, D. (2008). jModelTest: phylogenetic model averaging. Molecular Biology and Evolution, 25(7), 1253–1256.

Pulliandre, N., Lambert, A., Brouillet, S. and Achaz, G. (2011). ABGD, Automated Barcode Gap Discovery for primary species delineation. Molecular Ecology, 21, 1864–1877.

Rambaut, A., Drummond, A.J., Xie, D., Baele, G. and Suchard, M.A. (2018). Tracer v1.7, Available from http://tree.bio.ed.ac.uk/software/tracer/

Riesgo, A., Taboada, S. and Avila, C. (2015). Evolutionary patterns in Antarctic marine invertebrates: an update on molecular studies. Marine Genomics, 23, 1–13.

Rochebrune, A.T. and de Mabille, J. (1889). Mollusques. Mission Scientifique du Cap Horn 1882–1883. Tome 6 (Zoologie 2, part 8). Paris, Gauthiers-Villars, pls. 1–8, 11–12.

Ronquist, F., Huelsenbeck, J. and Teslenko, M. (2011). Draft MrBayes version 3.2 manual: tutorials and model summaries. Distributed with the software from http://brahms.biology.rochester.edu/software.html

Schiaparelli, S., Lörz, A.N. and Cattaneo-Vietti, R. (2006). Diversity and distribution of mollusc assemblages on the Victoria Land coast and the Balleny Islands, Ross Sea, Antarctica. Antarctic Science, 18(4), 615–631.

Schrödl, M., (1996). Nudibranchia y Sacoglossa de Chile: Morfología exterior y distribución. Gayana Zoología, 60, 17–62.

Schrödl, M., 2003. Sea slugs of Southern South America. 165.

Schrödl, M. (2009). Opisthobranchia - Sea Slugs. Marine Benthic Fauna of Chilean Patagonia, 505–542.

Schrödl, M., Jörger, K.M., Klussmann-Kolb, A. and Wilson, N.G. (2011). Bye bye “Opisthobranchia”! A review on the contribution of mesopsammic sea slugs to euthyneuran systematics. Thalassas, 27(2), 101–112.

Stamatakis, A., Hoover, P. and Rougemont, J. (2008). A rapid bootstrap algorithm for the RAxML web servers. Systematic Biology, 57(5), 758–771.

Stamatakis, A. (2014). RAxML version 8: a tool for phylogenetic analysis and post–analysis of large phylogenies. Bioinformatics, 30(9), 1312–1313.

Suchard, M.A., Lemey, P., Baele, G., Ayres, D.L., Drummond, A.J. and Rambaut, A. (2018). Bayesian phylogenetic and phylodynamic data integration using BEAST 1.10 Virus Evolution 4, vey016.

Talavera, G. and Castresana, J. (2007). Improvement of phylogenies after removing divergent and ambiguously aligned blocks from protein sequence alignments. Systematic Biology, 56(4), 564–577.

Thatje, S., Hillenbrand, C.D. and Larter, R. (2005). On the origin of Antarctic marine benthic community structure. Trends in Ecology and Evolution, 20(10), 534–540.

Thatje, S. (2012). Effects of capability for dispersal on the evolution of diversity in Antarctic benthos. Integrative and Comparative Biology, 52(4), 470–482.

Fujisawa, T. and Barraclough, T.G., (2013). Delimiting species using single-locus data and the Generalized Mixed Yule Coalescent approach: a revised method and evaluation on simulated data sets. Systematic Biology, 62(5), 707–724.

Wägele, H. (1995). The morphology and taxonomy of the Antarctic species of *Tritonia* Cuvier, 1797 (Nudibranchia: Dendronotoidea). Zoological Journal of the Linnean Society, 113(1), 21–46.

Wägele, H. and Willan, R.C. (2000). Phylogeny of the Nudibranchia. Zoological Journal of the Linnean Society, 130, 83–181.

Wägele, H., Ballesteros, M. and Avila, C., (2006). Defensive glandular structures in opisthobranch molluscs – from histology to ecology. Oceanography and Marine Biology, 44, 197.

Wägele, H., Klussmann-Kolb, A., Verbeek, E. and Schrödl, M. 2014. Flashback and foreshadowing – a review of the taxon Opisthobranchia. Organisms Diversity & Evolution, 14(1), 133–149.

Wilson, N.G., Schrödl, M. and Halanych, K.M. (2009). Ocean barriers and glaciation: evidence for explosive radiation of mitochondrial lineages in the Antarctic sea slug *Doris kerguelenensis* (Mollusca, Nudibranchia). Molecular Ecology, 18(5), 965–984.

Wilson, N.G., Maschek, J.A. and Baker, B.J. (2013). A species flock driven by predation? Secondary metabolites support diversification of slugs in Antarctica. PLoS ONE, 8(11), e80277.

WoRMS Editorial Board (2018). World Register of Marine Species. Available from http://www.marinespecies.org at VLIZ. Accessed 2018-02-27.

Vicente, N. and Arnaud, P.M. (1974). Invertébrés marins des XIIeme et XVeme expéditions Antarctiques Françaises en Terre Adélie. 12. Gastéropodes Opisthobranches. Tethys, 5, 531–548.

Yang, Z. (1996). Among-site rate variation and its impact on phylogenetic analyses. Trends in Ecology & Evolution, 11(9), 367–372.

