## Supplementary Material Table 1,2,3 for "Orange is the new white: taxonomic revision of Antarctic *Tritonia* species (Gastropoda: Nudibranchia)"

Running title: Antarctic *Tritonia* taxonomy

**Supplementary material 1.** Data of the specimens of *Tritonia* included in this study with including collection and geographical data.

| **Code** | **Cruise** | **Sample** | **Species** | **Preserved in** | **Station** | **Location** | **Latitude (S)** | **Longitude** | **Gear** | **Date** | **Depth (m)** |
| --- | --- | --- | --- | --- | --- | --- | --- | --- | --- | --- | --- |
| T01 | ANTXV3 | 67 | *T. challengeriana* | Frozen | 48/077 | North of Kapp Norvegia | 71° 10.2' | 12° 30.7' W | Agassiz trawl | 02/02/1998 | 433 |
| T02 | ANTXV3 | 78 | *T. challengeriana* | Frozen | 48/077 | North of Kapp Norvegia | 71° 10.2' | 12° 30.7' W | Agassiz trawl | 02/02/1998 | 433 |
| T03 | ANTXV3 | 79 | *T. challengeriana* | Frozen | 48/077 | North of Kapp Norvegia | 71° 10.2' | 12° 30.7' W | Agassiz trawl | 02/02/1998 | 433 |
| T04 | ANTXV3 | 80 | *T. challengeriana* | Frozen | 48/077 | North of Kapp Norvegia | 71° 10.2' | 12° 30.7' W | Agassiz trawl | 02/02/1998 | 433 |
| T05 | ANTXV3 | 118 | *T. challengeriana* | Frozen | 48/082 | Drescher inlet | 72° 50.5' | 19° 28' W | Bottom trawl | 03/02/1998 | 463 |
| T06 | ANTXV3 | 119 | *T. challengeriana* | Frozen | 48/082 | Drescher inlet | 72° 50.5' | 19° 28' W | Bottom trawl | 03/02/1998 | 463 |
| T07 | ANTXV3 | 127 | *T. challengeriana* | Frozen | 48/084 | Drescher inlet | 72° 50.5' | 19° 23' W | Bottom trawl | 03/02/1998 | 433 |
| T08 | ANTXV3 | 145 | *T. challengeriana* | Frozen | 48/097 | South of VestKapp | 73° 36.6' | 22°24.7' W | Bottom trawl | 05/02/1998 | 736 |
| T09 | ANTXV3 | 146 | *T. challengeriana* | Frozen | 48/097 | South of VestKapp | 73° 36.6' | 22°24.7' W | Bottom trawl | 05/02/1998 | 736 |
| T10 | ANTXV3 | 182 | *T. challengeriana* | Frozen | 48/150 | Halley Bay | 74°35.8' | 26°55.0' W | Bottom trawl | 11/02/1998 | 789 |
| T11 | ANTXV3 | 202 | *T. challengeriana* | Frozen | 48/189 | Kapp Norvegia | 71° 40.3' | 12°43.5' W | Agassiz trawl | 15/02/1998 | 244 |
| T12 | ANTXV3 | 267 | *T. challengeriana* | Frozen | 48/264 | Drescher Inlet | 72° 49.8' | 19°26.2'W | Agassiz trawl | 25/02/1998 | 471 |
| T13 | ANTXV3 | 268 | *T. challengeriana* | Frozen | 48/264 | Drescher Inlet | 72° 49.8' | 19°26.2' W | Agassiz trawl | 25/02/1998 | 471 |
| T14.1 | ANTXXI-2 | 22 | *T. challengeriana* | Frozen | PS65/019-1 | Bouvet Island | 54° 30.01' | 3° 13.97' E | Agassiz trawl | 24/11/2003 | 259.7 |
| T14.2 | ANTXXI-2 | 22 | *T. challengeriana* | Frozen | PS65/019-1 | Bouvet Island | 54° 30.01' | 3° 13.97' E | Agassiz trawl | 24/11/2003 | 259.7 |
| T14.3 | ANTXXI-2 | 22 | *T. challengeriana* | Frozen | PS65/019-1 | Bouvet Island | 54° 30.01' | 3° 13.97' E | Agassiz trawl | 24/11/2003 | 259.7 |
| T14.4 | ANTXXI-2 | 22 | *T. challengeriana* | Frozen | PS65/019-1 | Bouvet Island | 54° 30.01' | 3° 13.97' E | Agassiz trawl | 24/11/2003 | 259.7 |
| T14.5 | ANTXXI-2 | 22 | *T. challengeriana* | Frozen | PS65/019-1 | Bouvet Island | 54° 30.01' | 3° 13.97' E | Agassiz trawl | 24/11/2003 | 259.7 |
| T14.6 | ANTXXI-2 | 22 | *T. challengeriana* | Frozen | PS65/019-1 | Bouvet Island | 54° 30.01' | 3° 13.97' E | Agassiz trawl | 24/11/2003 | 259.7 |
| T14.7 | ANTXXI-2 | 22 | *T. challengeriana* | Frozen | PS65/019-1 | Bouvet Island | 54° 30.01' | 3° 13.97' E | Agassiz trawl | 24/11/2003 | 259.7 |
| T14.8 | ANTXXI-2 | 22 | *T. challengeriana* | Frozen | PS65/019-1 | Bouvet Island | 54° 30.01' | 3° 13.97' E | Agassiz trawl | 24/11/2003 | 259.7 |
| T15.1 | ANTXXI-2 | 61 | *T. dantarti* | 70% Ethanol | PS65/028-1 | Bouvet Island | 54° 22.49' | 3° 17.58' E | Agassiz trawl | 25/11/1999 | 130 |
| T15.2 | ANTXXI-2 | 61 | *T. dantarti* | 70% Ethanol | PS65/028-1 | Bouvet Island | 54° 22.49' | 3° 17.58' E | Agassiz trawl | 25/11/1999 | 130 |
| T15.3 | ANTXXI-2 | 61 | *T. dantarti* | 70% Ethanol | PS65/028-1 | Bouvet Island | 54° 22.49' | 3° 17.58' E | Agassiz trawl | 25/11/1999 | 130 |
| T15.4 | ANTXXI-2 | 61 | *T. dantarti* | 70% Ethanol | PS65/028-1 | Bouvet Island | 54° 22.49' | 3° 17.58' E | Agassiz trawl | 25/11/1999 | 130 |
| T15.5 | ANTXXI-2 | 61 | *T. dantarti* | 70% Ethanol | PS65/028-1 | Bouvet Island | 54° 22.49' | 3° 17.58' E | Agassiz trawl | 25/11/1999 | 130 |
| T16 | ANTXXI-2 | 62 | *T. dantarti* | 96% Ethanol | PS65/028-1 | Bouvet Island | 54° 22.49' | 3° 17.58' E | Agassiz trawl | 25/11/2000 | 131 |
| T17.1 | ANTXXI-2 | 63 | *T. dantarti* | Frozen | PS65/028-1 | Bouvet Island | 54° 22.49' | 3° 17.58' E | Agassiz trawl | 25/11/2001 | 132 |
| T17.2 | ANTXXI-2 | 63 | *T. dantarti* | Frozen | PS65/028-1 | Bouvet Island | 54° 22.49' | 3° 17.58' E | Agassiz trawl | 25/11/2001 | 132 |
| T17.3 | ANTXXI-2 | 63 | *T. dantarti* | Frozen | PS65/028-1 | Bouvet Island | 54° 22.49' | 3° 17.58' E | Agassiz trawl | 25/11/2001 | 132 |
| T17.4 | ANTXXI-2 | 63 | *T. dantarti* | Frozen | PS65/028-1 | Bouvet Island | 54° 22.49' | 3° 17.58' E | Agassiz trawl | 25/11/2001 | 132 |
| T18.1 | ANTXXI-2 | 64 | *T. dantarti* | Karnovsky | PS65/028-1 | Bouvet Island | 54° 22.49' | 3° 17.58' E | Agassiz trawl | 25/11/2002 | 133 |
| T18.2 | ANTXXI-2 | 64 | *T. dantarti* | Karnovsky | PS65/028-1 | Bouvet Island | 54° 22.49' | 3° 17.58' E | Agassiz trawl | 25/11/2002 | 133 |
| T19.1 | ANTXXI-2 | 84 | *T. challengeriana* | Frozen | PS65/029-1 | Bouvet Island | 54° 31.59' | 3° 13.05' E | Agassiz trawl | 25/11/2003 | 376.8 |
| T19.2 | ANTXXI-2 | 84 | *T. challengeriana* | Frozen | PS65/029-1 | Bouvet Island | 54° 31.59' | 3° 13.05' E | Agassiz trawl | 25/11/2003 | 376.8 |
| T19.3 | ANTXXI-2 | 84 | *T. challengeriana* | Frozen | PS65/029-1 | Bouvet Island | 54° 31.59' | 3° 13.05' E | Agassiz trawl | 25/11/2003 | 376.8 |
| T19.4 | ANTXXI-2 | 84 | *T. challengeriana* | Frozen | PS65/029-1 | Bouvet Island | 54° 31.59' | 3° 13.05' E | Agassiz trawl | 25/11/2003 | 376.8 |
| T19.5 | ANTXXI-2 | 84 | *T. challengeriana* | Frozen | PS65/029-1 | Bouvet Island | 54° 31.59' | 3° 13.05' E | Agassiz trawl | 25/11/2003 | 376.8 |
| T19.6 | ANTXXI-2 | 84 | *T. challengeriana* | Frozen | PS65/029-1 | Bouvet Island | 54° 31.59' | 3° 13.05' E | Agassiz trawl | 25/11/2003 | 376.8 |
| T20 | ANTXXI-2 | 103 | *T.* *dantarti* | 70% Ethanol | PS65/028-1 | Bouvet Island | 54° 22.49' | 3° 17.58' E | Agassiz trawl | 25/11/2003 | 134 |
| T21 | ANTXXI-2 | 104 | *T. challengeriana* | 70% Ethanol | PS65/029-1 | Bouvet Island | 54° 31.59' | 3° 13.05' E | Agassiz trawl | 25/11/2003 | 376.8 |
| T22 | ANTXXI-2 | 107 | *T. challengeriana* | 96% Ethanol | PS65/029-1 | Bouvet Island | 54° 31.59' | 3° 13.05' E | Agassiz trawl | 25/11/2003 | 376.8 |
| T23 | ANTXXI-2 | 117 | *T. challengeriana* | Frozen | PS65/039-1 | North of Kapp Norvegia | 71° 06.30' | 11° 32.04' W | Agassiz trawl | 05/12/2003 | 175.2 |
| T24 | ANTXXI-2 | 332 | *T. challengeriana* | Frozen | PS65/121-1 | North of Kapp Norvegia | 70° 50.08' | 10° 34.76' W | Agassiz trawl | 11/12/2003 | 274 |
| T25 | ANTXXI-2 | 779 | *T. challengeriana* | 70% Ethanol | PS65/251-1 | North of Kapp Norvegia | 71° 07.34' | 11° 27.80' W | Rauschert dredge | 23/12/2003 | 145.6 |
| T26 | ANTXXI-2 | 1020 | *T. challengeriana* | Frozen | PS65/276-1 | North of Kapp Norvegia | 71° 06.44' | 11° 27.76' W | Agassiz trawl | 28/12/2003 | 277.2 |
| T27.1 | ANTXXI-2 | 1030 | *T. challengeriana* | Frozen | PS65/276-1 | North of Kapp Norvegia | 71° 06.44' | 11° 27.76' W | Agassiz trawl | 28/12/2003 | 277.2 |
| T27.2 | ANTXXI-2 | 1030 | *T. challengeriana* | Frozen | PS65/276-1 | North of Kapp Norvegia | 71° 06.44' | 11° 27.76' W | Agassiz trawl | 28/12/2003 | 277.2 |
| T28 | ANTXXI-2 | 1161 | *T. challengeriana* | 10% Formaline | PS65/292-1 | Drescher Inlet | 72° 51.43' | 19° 38.62' W | Bottom trawl | 31/12/2003 | 597.6 |
| T29 | ANTXXI-2 | 1225 | *T. challengeriana* | 10% Formaline | PS65/308-1 | Drescher Inlet | 72° 50.18' | 19° 35.94' W | Rauschert dredge | 02/01/2004 | 622 |

**Supplementary material 2.** Data of samples of the species included in the phylogenetic analyses, with Voucher numbers and access codes (COI, 16S, H3) from GenBank. Our samples include the 16 specimens used for molecular analysis.

| Code | Species | Voucher nº | COI | 16S | H3 | Locality | Reference |
| --- | --- | --- | --- | --- | --- | --- | --- |
| BANG | *Bornella anguilla* | CASIZ 191601 | KP871636 | KP871683 | KP871659 | – | Mahguib et al. (2015) |
| BHER | *Bornella hermanni* | CASIZ 175743 | HM162705 | HM162625 | HM162531 | Malaysia: Tokong Kamundi | Pola and Gosliner (2010) |
| BJOH | *Bornella johnsonorum* | CASIZ 175406 | HM162704 | HM162624 | HM162530 | Marshall Islands: Kwajalein Atoll | Pola and Gosliner (2010) |
| BSTE | *Bornella stellifer* | CASIZ 167989 | HM162703 | HM162623 | HM162529 | USA: Hawaii, Lanai | Pola and Gosliner (2010) |
| BVAL | *Bornella valdae* | CASIZ 176832 | HM162706 | HM162626 | HM162532 | South Africa: Durban | Pola and Gosliner (2010) |
| CGRA | *Curnon granulosa* | – | GQ292060 | – | – | Antarctica: Ross sea | Shields et al. (2009) |
| DALB | *Dirona albolineata* | 10BCMOL-00344 | KF643932 | – | – | – | Layton et al. (2014) |
| DPIC | *Dirona picta* | – | DQ026831 | – | – | – | Heminez et al. (2005) |
| LMIL | *Leminda millecra* | CASIZ 176348 | HM162745 | HM162669 | HM162578 | S.A.: Western Cape Province | Pola and Gosliner (2010) |
| MARB | *Marionia arborescens* | CAS:177735 | KP226855 | KP226859 | KP226857 | Philippines | Mahguib et al. (2015) |
| MBLA | *Marionia blainvillea* | CASIZ 176812 | HM162721 | HM162645 | HM162553 | Portugal: Azores | Pola and Gosliner (2010) |
| MDIS | *Marionia distincta* | CAS:110364 | KP226856 | KP226860 | KP226858 | – | Mahguib et al. (2015) |
| MELO | *Marionia elongoviridis* | CASIZ 173308 | HM162724 | – | HM162556 | Philippines: Panglao | Pola and Gosliner (2010) |
| MLEV | *Marionia levis* | CASIZ 173454 | HM162723 | HM162647 | HM162555 | Madagascar: Kalakajoro | Pola and Gosliner (2010) |
| MSP5 | *Marionia sp. 5* | CASIZ 177513 | HM162727 | HM162650 | HM162559 | Philippines: Batangas | Pola and Gosliner (2010) |
| T01 | *Tritonia challengeriana* | – | – | MN648383 | MN651111 | Weddell Sea | This study |
| T02 | *Tritonia challengeriana* | – | – | MN648384 | MN651112 | Weddell Sea | This study |
| T03 | *Tritonia challengeriana* | – | MN651126 | MN648385 | MN651113 | Weddell Sea | This study |
| T05 | *Tritonia challengeriana* | – | – | MN648386 | MN651114 | Weddell Sea | This study |
| T06 | *Tritonia challengeriana* | – | MN651127 | MN648386 | MN651115 | Weddell Sea | This study |
| T07 | *Tritonia challengeriana* | – | MN651128 | MN648387 | MN651116 | Weddell Sea | This study |
| T08 | *Tritonia challengeriana* | – | MN651129 | MN648388 | MN651117 | Weddell Sea | This study |
| T09 | *Tritonia challengeriana* | – | MN651130 | MN648389 | MN651118 | Weddell Sea | This study |
| T10 | *Tritonia challengeriana* | – | MN651131 | MN648390 | MN651119 | Weddell Sea | This study |
| T11 | *Tritonia challengeriana* | – | – | MN648391 | MN651120 | Weddell Sea | This study |
| T12 | *Tritonia challengeriana* | – | MN651132 | MN648392 | MN651121 | Weddell Sea | This study |
| T13 | *Tritonia challengeriana* | – | MN651133 | MN648393 | MN651122 | Weddell Sea | This study |
| T14.3 | *Tritonia dantarti* | – | MN651134 | MN648394 | – | Bouvet Island | This study |
| T17.1 | *Tritonia dantarti* | – | MN651135 | – | MN651123 | Bouvet Island | This study |
| T17.2 | *Tritonia dantarti* | – | MN651136 | MN648395 | MN651124 | Bouvet Island | This study |
| T22 | *Tritonia dantarti* | – | – | MN648396 | MN651125 | Bouvet Island | This study |
| TANT | *Tritonia antarctica* | CASIZ 171177 | HM162718 | HM162643 | HM162550 | Bouvet Island | Pola and Gosliner (2010) |
| TBEL1 | *Tritoniella belli* | – | GU227111 | GU227002 | – | McMurdo Sound, Ross Sea | Heimeier et al. (2010) |
| TBEL2 | *Tritoniella belli* | – | GQ292056 | – | – | Ross Sea | Shields et al. (2009) |
| TCHA | *Tritonia challengeriana* | – | GQ292052 | – | – | Ross Sea | Shields et al. (2009) |
| TFES1 | *Tritonia festiva* | CASIZ 186478 | KP153291 | KP153258 | KP153324 | – | Hulett et al. (2014) |
| TFES2 | *Tritonia festiva* | CASIZ 174491 | HM162719 | – | HM162551 | Oregon, Coos County | Pola and Gosliner (2010) |
| THAM | *Tritonia hamnerorum* | CASIZ 181090 | KP153293 | KP153260 | KP153326 | – | Hulett et al. (2014) |
| THOM | *Tritonia hombergii* | MT09685 | KR084797 | – | – | North Sea | Barco et al. (2015) |
| TNIL1 | *Tritonia nilsodhneri* | CASIZ 176219 | HM162716 | HM162641 | HM162548 | Cape Province, False Bay | Pola and Gosliner (2010) |
| TNIL2 | *Tritonia nilsodhneri* | CASIZ 176222 | KP871653 | KP871702 | KP871677 | – | Mahguib et al. (2015) |
| TNIL3 | *Tritonia nilsodhneri* | CASIZ 176218B | KP153295 | KP153262 | KP153328 | South Africa | Pola and Gosliner (2015) |
| TNIL4 | *Tritonia nilsodhneri* | CASIZ 176220 | KP940454 | KP940449 | KP940459 | – | Hulett et al. (2014) |
| TNIL5 | *Tritonia nilsodhneri* | CASIZ 176218A | KP153294 | KP153261 | KP153327 | – | Hulett et al. (2014) |
| TPIC | *Tritonia pickensi* | CASIZ 175718 | HM162717 | HM162642 | HM162549 | Costa Rica: Islas Catalinas | Pola and Gosliner (2010) |
| TPLE4 | *Tritonia plebeia* | ZMMU Op-572 | KX788134 | KX788122 | – | Norway | Korshunova et al. (2016) |
| TPLE1 | *Tritonia plebeia* | MT09662 | KR084895 | – | – | North Sea | Barco et al. (2015) |
| TPLE2 | *Tritonia plebeia* | MT09661 | KR084842 | – | – | North Sea | Barco et al. (2015) |
| TPLE3 | *Tritonia plebeia* | MT09675 | KR084634 | – | – | North Sea | Barco et al. (2015) |
| TSP3 | *Tritonia sp. 3* | CASIZ 177523 | HM162731 | HM162654 | HM162563 | Luzon, Twin Rocks | Pola and Gosliner (2010) |
| TSP4 | *Tritonia sp. 4* | CASIZ 177668 | HM162732 | HM162655 | HM162564 | Philippines: Batangas, Anilao | Pola and Gosliner (2010) |
| TSP5 | *Tritonia sp. 5* | CASIZ 173748 | KP153303 | KP153270 | KP153336 | – | Hulett et al. (2014) |
| TSP6 | *Tritonia sp. 6* | CASIZ 190807 | KP153301 | KP153268 | KP153334 | – | Hulett et al. (2014) |
| TSP7 | *Tritonia sp. 7* | CASIZ 191401 | KP153300 | KP153267 | KP153333 | – | Hulett et al. (2014) |
| TSPF | *Tritonia sp. F* | CASIZ 179495 | HM162720 | HM162644 | HM162552 | Batangas, Luzon, Tingloy, Bethlem | Pola and Gosliner (2010) |
| TSPG | *Tritonia sp. G* | CASIZ 176233 | HM162730 | HM162653 | HM162562 | Cape Province, False Bay | Pola and Gosliner (2010) |
| TSTR1 | *Tritonia striata* | BAU2696 | LT596541 | LT596543 | LT615408 | Italy, Grosseto, Le formiche Is | Furfaro et al. (2016) |
| TSTR2 | *Tritonia striata* | BAU2695 | LT596540 | LT596540 | LT615407 | Italy, Grosseto, Le formiche Is | Furfaro et al. (2016) |

**Supplementary material 3.** Best-fit models and parameters calculated in jModeltest and Gblocks.

| Parameters | COI | 16S | 16S Relaxed | 16S Stringent | H3 |
| --- | --- | --- | --- | --- | --- |
| Nº specimens | 53 | 45 | 45 | 45 | 47 |
| Nº Characters | 601 | 486 | 439 | 358 | 328 |
| Best-fit model | TrN+I+G | TPM3uf+I+G | TPM3uf+I+G | GTR+I+G | TIM1+I+G |
| Frequency A | 0.276 | 0.342 | 0.348 | 0.32 | 0.253 |
| Frequency C | 0.097 | 0.12 | 0.121 | 0.134 | 0.323 |
| Frequency G | 0.2 | 0.187 | 0.121 | 0.227 | 0.262 |
| Frequency T | 0.428 | 0.351 | 0.353 | 0.319 | 0.162 |
| Γ shape (g) | 0.372 | 0.449 | 0.443 | 0.428 | 0.396 |
| Proportion of invariant sites (i) | 0.484 | 0.330 | 0.370 | 0.379 | 0.365 |
| R-matrix [A–C] | 1 | 0.511 | 0.528 | 2.904 | 1 |
| R-matrix [A–G] | 15.729 | 3.373 | 3.624 | 6.325 | 52.879 |
| R-matrix [A–T] | 1 | 1 | 1 | 2.904 | 14.219 |
| R-matrix [C–G] | 1 | 0.511 | 0.528 | 1 | 14.219 |
| R-matrix [C–T] | 52.466 | 3.373 | 3.625 | 11.893 | 129.314 |
| R-matrix [G–T] | 1 | 1 | 1 | 1 | 10 |
| Min. Nº seq. Conserved pos. | – | – | 24 | 24 | – |
| Min. Nº seq. Flank pos. | – | – | 24 | 39 | – |
| Max. Nº contig. Non-conserved pos. | – | – | 8 | 4 | – |
| Min. Length block | – | – | 5 | 10 | – |
| Allowed gap pos. | – | – | Half | None | – |

**Supplementary material 4**. GMYC analysis results suggesting valid species groups given our tree topology.

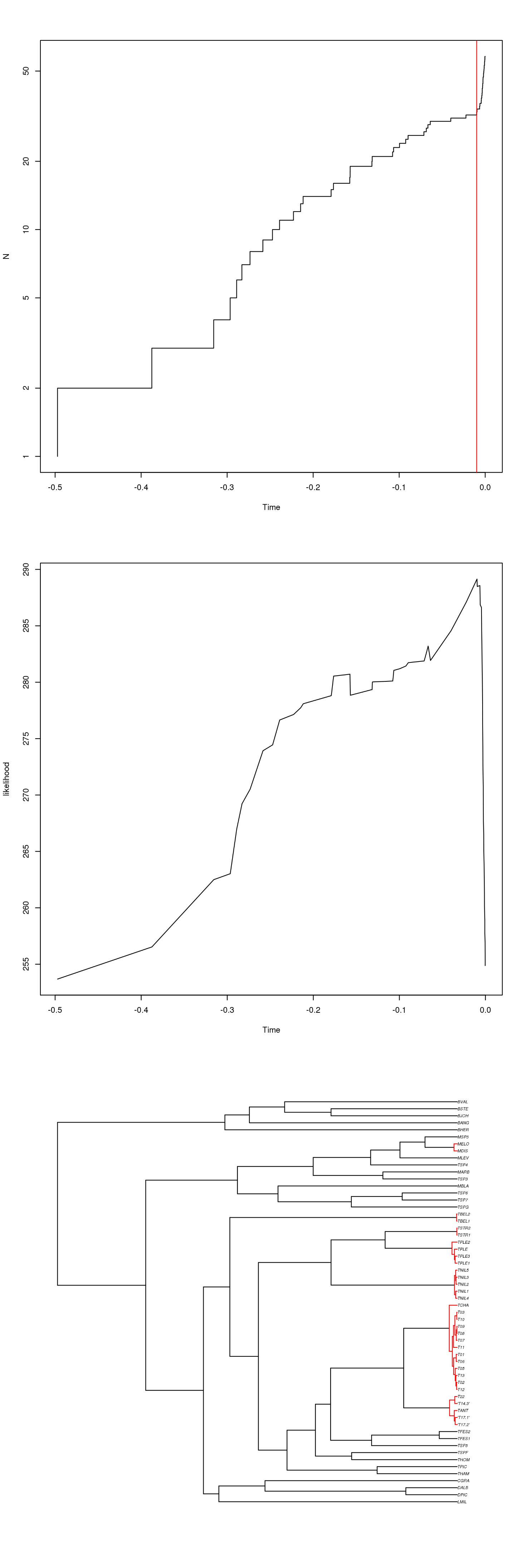

**References**

Barco, A., Raupach, M.J., Laakmann, S., Neumann, H. and Knebelsberger, T., 2016. Identification of North Sea molluscs with DNA barcoding. *Molecular ecology resources*, *16*(1), pp.288-297.

Furfaro, G., Mariottini, P., Modica, M.V., Trainito, E., Doneddu, M. and Oliverio, M. (2016). Sympatric sibling species: the case of *Caloria elegans* and *Facelina quatrefagesi* (Gastropoda: Nudibranchia). *Scientia Marina*, 80(4), 511–520.

Heimeier, D., Lavery, S. and Sewell, M.A., 2010. Using DNA barcoding and phylogenetics to identify Antarctic invertebrate larvae: Lessons from a large scale study. *Marine genomics*, *3*(3-4), pp.165-177.

Heminez,L.A., Pence,W.E. and Mason,D.E. Unpublished

Hulett, R.E., and Gosliner, T.M. (2014). Rooting for clarity: a phylogenetic reconstruction of the nudibranch family Tritoniidae. *Integrative and Comparative Biology*, 54, E290–E290.

Korshunova, T., Sanamyan, N., Zimina, O., Fletcher, K. and Martynov, A. (2016). Two new species and a remarkable record of the genus *Dendronotu*s from the North Pacific and Arctic oceans (Nudibranchia). *ZooKeys,* (630), 19.

Layton, K.K., Martel, A.L. and Hebert, P.D. (2014). Patterns of DNA barcode variation in Canadian marine molluscs. *PLoS ONE*, 9(4), p.e95003.

Mahguib, J. and Valdés, Á. (2015). Molecular investigation of the phylogenetic position of the polar nudibranch *Doridoxa* (Mollusca, Gastropoda, Heterobranchia). *Polar Biology*, 38(9), 1369–1377.

Pola, M. and Gosliner, T.M. (2010). The first molecular phylogeny of cladobranchian opisthobranchs (Mollusca, Gastropoda, Nudibranchia). *Molecular Phylogenetics and Evolution*, 56(3), 931–941.

Pola, M. and Gosliner, T.M. (2015). A new large and colourful species of the genus *Doto* (Nudibranchia: Dotidae) from South Africa. *Journal of Natural History*, 49(41-42), pp.2465–2481.

Shields, C.C. (2009). Nudibranchs of the Ross Sea, Antarctica: Phylogeny, diversity, and divergence. Master thesis: Clemson University
